# Characterization of L-serine deaminases, SdaA (PA2448) and SdaB (PA5379), and their potential role in *Pseudomonas aeruginosa* pathogenesis

**DOI:** 10.1101/394957

**Authors:** Sixto M. Leal, Elaine Newman, Kalai Mathee

**Affiliations:** Department of Biological Sciences, College of Arts Sciences and Education, Florida International University, Miami, United States of America; Department of Biological Sciences, Concordia University, Montreal, Canada; Department of Molecular Microbiology and Infectious Diseases, Herbert Wertheim College of Medicine, Florida International University, Miami, United States of America; Biomolecular Sciences Institute, Florida International University, Miami, United States of America; Department of Human and Molecular Genetics, Herbert Wertheim College of Medicine, Florida International University, Miami, United States of America; Case Western Reserve University, United States of America

**Keywords:** Serine Catabolism, Central Metabolism, TCA Cycle, Pyruvate, Leucine Responsive Regulatory Protein (LRP), One Carbon Metabolism

## Abstract

Regardless of the site of infectivity, all pathogens require high energetic influxes. This energy is required to counterattack the host immune system and in the absence the bacterial infections are easily cleared by the immune system. This study is an investigation into one highly bioenergetic pathway in *Pseudomonas aeruginosa* involving the amino acid L-serine and the enzyme L-serine deaminase (L-SD). *P. aeruginosa* is an opportunistic pathogen causing infections in patients with compromised immune systems as well as patients with cystic fibrosis. L-SD has been linked directly to the pathogenicity of several organisms including but not limited to *Campylobacter jejuni, Mycobacterium bovis*, *Streptococcus pyogenes*, and *Yersinia pestis*. We hypothesized that *P. aeruginosa* L-SD is likely to be critical for its virulence. The genome sequence analysis revealed the presence of two L-SD homologs encoded by *sdaA* and *sdaB.* We analyzed the ability of *P. aeruginosa* to utilize serine and the role of SdaA and SdaB in serine deamination by comparing mutant strains of *sdaA* (PAO*sdaA*) and *sdaB* (PAO*sdaB*) with their isogenic parent *P. aeruginosa* PAO1. We demonstrate that *P. aeruginosa* is unable to use serine as a sole carbon source. However, serine utilization is enhanced in the presence of glycine. Both SdaA and SdaB contribute to L-serine deamination, 34 % and 66 %, respectively. Glycine was also shown to increase the L-SD activity especially from SdaB. Glycine-dependent induction requires the inducer serine. The L-SD activity from both SdaA and SdaB is inhibited by the amino acid L-leucine. These results suggest that *P. aeruginosa* L-SD is quite different from the characterized *E. coli* L-SD that is glycine-independent but leucine-dependent for activation. Growth mutants able to use serine as sole carbon source were isolated. In addition, suicide vectors were constructed which allow for selective mutation of the *sdaA* and *sdaB* genes on any *P. aeruginosa* strain of interest. Future studies with a double mutant will reveal the importance of these genes for pathogenicity.

## INTRODUCTION

Amino acids can be used as a carbon source. Most of the amino acids are converted to pyruvate that enters the Tricarboxylic Acid Cycle (TCA). In the absence of glucose, microorganisms preferentially utilize L-serine as a carbon source (Zinser & Kolter, 2000; Zinser & Kolter, 2004; Prüß, 1994). L-serine is catabolized to pyruvate after a single enzymatic reaction catalyzed by L-serine deaminase (L-SD) (Prüß, 1994). The pyruvate is converted to acetyl CoA, which is oxidized via the TCA cycle to carbon dioxide and free energy (Figure 1) L-serine also serves as an anabolic substrate for vital functions within the cell. It serves as a monomer for proteins, plays a significant role in phospholipid biosynthesis and acts as a substrate for the enzymatic synthesis of glycine, cysteine, and tryptophan (Stauffer, 1996). Glycine, in turn, is a precursor for purines and heme groups involved in essential nucleic acid formation and redox reactions, respectively (Stauffer, 1996). Thus, L-serine has a central role in bacterial metabolism. Fifteen % of the carbon assimilation entering *Escherichia coli* metabolism from extracellular glucose stores is at one point or another incorporated into a molecule of serine (Pizer & Potochny, 1964).

**Figure 1.**
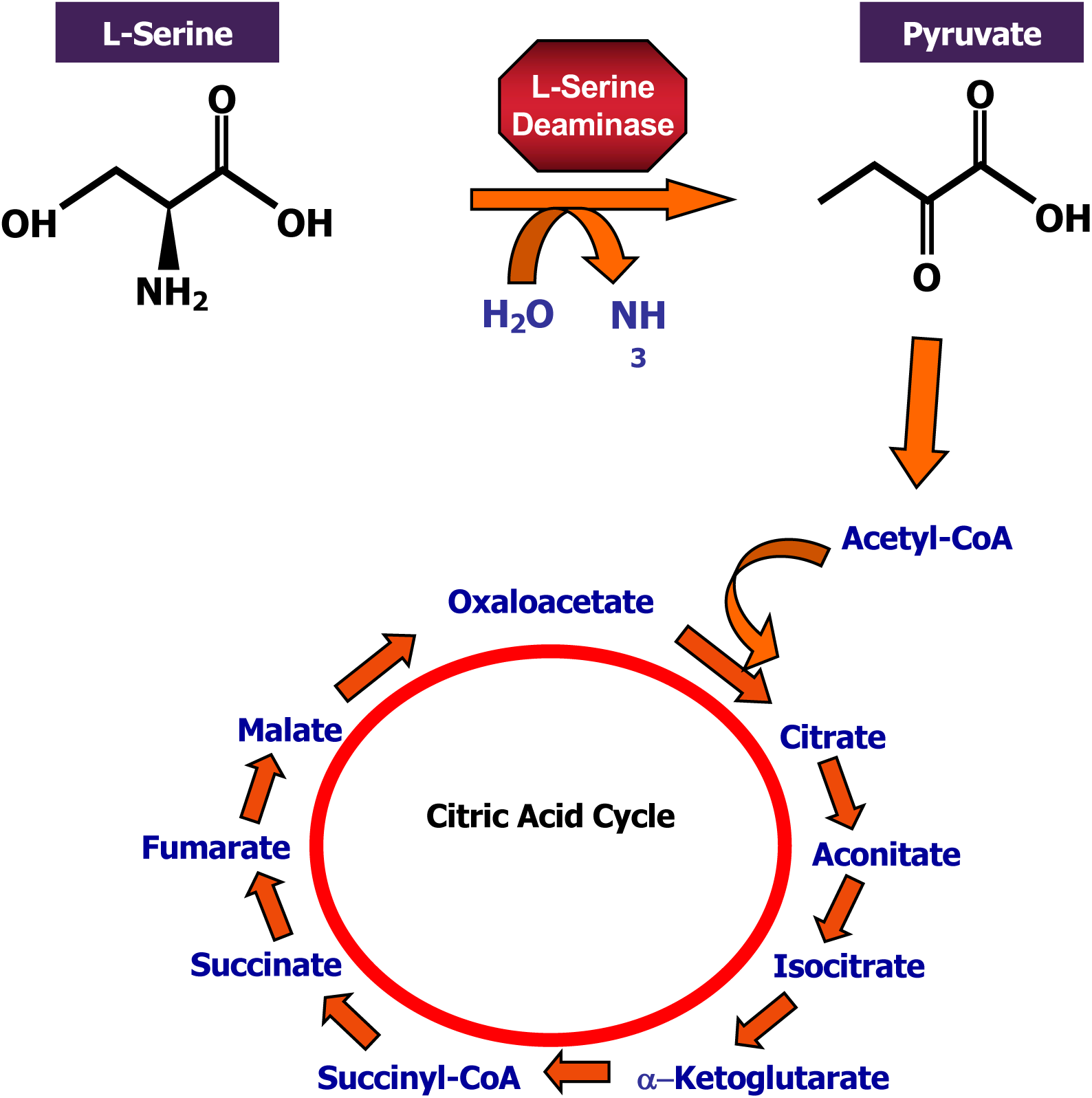
L-Serine catabolism. L-serine is converted to pyruvate via a one step reaction catalyzed by L-serine deaminase (L-SD). One molecule of H_2_O is absorbed and one molecule of NH_3_ is released during the process. Pyruvate then enters the citric acid cycle yielding one GTP, four NADH_2_, and one FADH_2_ molecule.

Despite L-serine’s known central role in bacterial metabolism, most organisms do not utilize L-serine as the sole carbon source (McFall, 1996; Mendz & Hazell, 1995; Vining & Magasanik, 1981). When in complex media, however, L-serine is the first amino acid catabolized (Prüß, 1994). Specifically, serine degradation is greatly enhanced in the presence of small quantities of other carbon sources such as glucose, glycine, leucine, isoleucine, or threonine (Ogawa *et al.*, 1997). Under these conditions there are two reported phases of serine utilization during the cell growth cycle (Mendz & Hazell, 1995; Novak, 2000; Velayudhan & Kelly, 2002). In the first phase most of the serine is metabolized to pyruvate with the subsequent energy and intermediates contributing to the growth of the bacterium (Prüß, 1994; Netzer *et al.*, 2004). In the second phase serine is still consumed, but no further growth is observed (Prüß, 1994; Netzer *et al.*, 2004). The ability to utilize serine in this second phase of growth, allows for preferential survival by these microorganisms (Zinser & Kolter, 2000).

L-serine degradation is preferred over that of glucose in *E. coli* (McFall, 1996). Biochemically, serine degradation is less complex, involving fewer steps than glucose catabolism. Glucose is broken down via glycolysis (Figure 2) into two molecules of pyruvate, two adenosine triphosphates (ATP), and two nicotinamide dinucleotide (NADH) molecules by substrate level phosphorylation of adenosine diphosphate (ADP). This metabolic process uses 10 enzymes, each of which is produced by the transcription of their corresponding genes and translation of the corresponding protein (Stover *et al.*, 2000). The net outcome of glycolysis is bio energetically favorable. However, in comparison to serine catabolism, the breakdown of serine is more energetically favorable (Figure 2). Serine utilization by the organism requires transcription of one gene and translation of one protein. As compared to glycolysis, serine catabolism bypasses ten reactions which yield 38 ATP molecules post oxidative phosphorylation, for one reaction that directly feeds into the Krebs cycle leading to the production of 15 ATPs per pyruvate molecule (Figure 2). There is a 15:1 ATP to enzyme ratio for L-serine catabolism as compared to a 3.8:1 in glycolysis. This 4 fold increase in bioenergetic efficiency could explain the observed contribution of L-serine catabolism to bacterial pathogenicity (Velayudhan *et al.*, 2004; Chen *et al.*, 2003).

**Figure 2.**
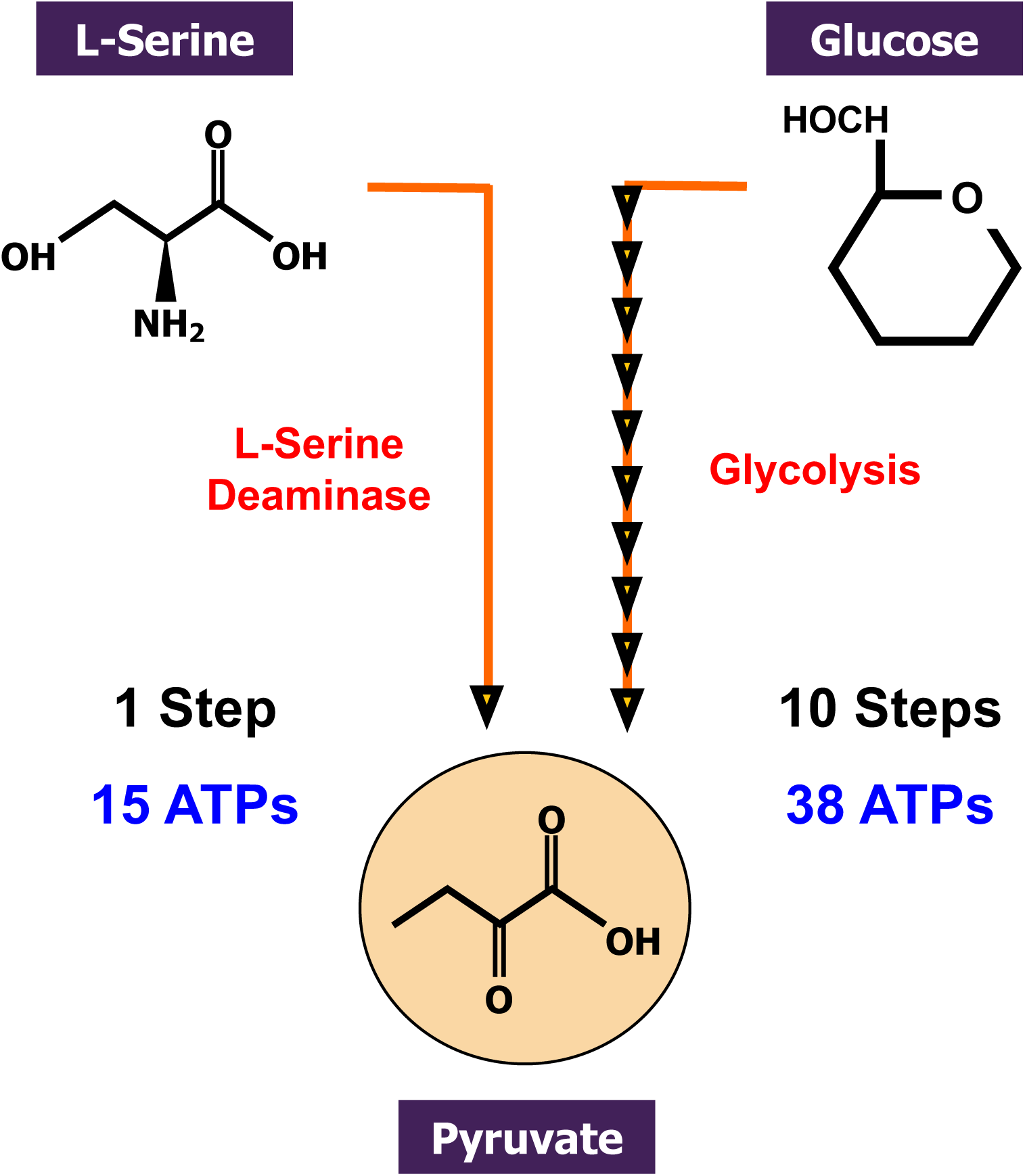
Comparison of L-serine deamination with glycolysis. In comparison to glycolysis which involves ten enzymes ultimately leading to the production of 38 ATPs, L-serine catabolism only requires one enzyme, L-serine deaminase (L-SD) ultimately leading to the production of 15 ATPs. There is a 15:1 ATP to enzyme ratio for L-serine catabolism as compared to a 3.8:1 ATP to enzyme ratio in glycolysis.

The enzyme that catalyzes the conversion of L-serine to pyruvate and ammonia is L-Serine Deaminase (Pardee & Prestidge, 1955; Simmonds & Miller, 1957; Pardee & Prestidge, 1955). *E. coli* harbors two L-SDs, LSD2 and LSD3 encoded by the genes *sdaB, and tdcG. E. coli* also harbors a third L-serine delaminating enzyme known as L-serine ammonia lyase encoded by *sdaA*, with affinity for both amino acids, threonine and serine (Keseler *et al.*, 2005). In *E. coli* L-SD is translated in an inactive form and activated post translationally via synthesis of a 4Fe4S cluster by the gene products of the *isc* operon (Cicchillo *et al.*, 2004). Prokaryotic L-SD differs in its catalytic mechanism from known eukaryotic L-SDs. While eukaryotic L-SD carries out the deamination step via the amino-transferring coenzyme pyridoxal phosphate (Ogawa *et al.*, 1989), the prokaryotic homolog uses an iron sulfur center for catalysis (4Fe-4S) (Cicchillo *et al.*, 2004). In *E. coli* the regulation of genes encoding L-SD are under control of the **L**eucine **R**esponsive Regulatory **P**rotein (Lrp) (Newman *et al.*, 1985).

In order for a pathogen to successfully colonize its host it must acquire energy from the host. This is most efficiently achieved via secretion of proteases to break down neighboring host proteins to smaller peptides and or individual amino acids. Given the selective advantage conferred on organisms with high L-SD activity *in vitro*, the ability to utilize L-serine released by proteolytic cleavage *in vivo* should confer a selective advantage to these pathogenic organisms as well. Indeed, L-SD has been implicated in the pathogenicity of several organisms, including *Campylobacter jejuni, Mycobacterium bovis, and Streptococcus pyogenes.* In *C. jejuni,* isogenic L-SD mutants failed to colonize the avian gut, while all of the wild type strains inoculated into newborn chickens were able to colonize the cecum (Velayudhan *et al.*, 2004). Thus it appears that L-serine degradation is critical for *C. jejuni* pathogenicity.

Any invading organism must break down host metabolites and convert them to free energy, which it can use to sustain its growth in the host. Without the ability to harness energy from these catabolites, such pathogens cannot persist against the massive attacks by the host immune system. The inability to degrade L-serine has been linked to the failure of the live-attenuated vaccine for *M. tuberculosis* to persist in its host (Chen *et al.*, 2003). The Bacille Calmette-Guerin (BCG) vaccine, administered worldwide except in the United States, was shown to have an intact gene encoding L-SD, however expression was undetectable (Chen *et al.*, 2003). Lack of L-SD expression led to the accumulation of serine in the cell, which was then shown to inhibit glutamine synthase activity (Chen *et al.*, 2003). Thus it has been proposed that the low L-SD expression resulted in the failure of the live vaccine to establish a persistent infection (Chen *et al.*, 2003).

Given the selective pressure of host defenses, many pathogenic organisms have devised virulence factors that have allowed them to persist under attack. The production of these virulence factors is often excessive, such as the excess polysaccharide coating produced by many organisms to resist phagocytosis, complement, and opsonization. The production of virulence factors requires energy, and thus some organisms regulate virulence factor gene expression with metabolic gene expression. For example, *Streptococcus pyogenes* produces the Rgg protein, which serves as a transcriptional regulator that switches virulence genes on and off as the organism prepares to invade its host (Chaussee *et al.*, 2003). Besides the expression of virulence factors, it also regulates the expression of L-SD encoding genes (Chaussee *et al.*, 2003). Thus, a virulence factor gene regulator manipulates the expression of a metabolic gene when the bacterium is preparing to infect. This finding adds credence to the hypothesized contribution of L-SD expression to bacterial pathogenicity.

The role of L-SD in the pathogenicity of *P. aeruginosa* has not been characterized. This Gram negative facultative aerobe is ubiquitous in nature and is opportunistically pathogenic to burn patients, immunocompromised individuals, and people suffering from cystic fibrosis (Lyczak *et al.*, 2002). *P. aeruginosa*’s mode of growth protects the organism from antibiotics, as well as antibodies, complement, and opsonization (Govan & Harris, 1986). Upon selection by host’s defenses, the bacterial population switches from a non-mucoid to a mucoid form (Mathee *et al.*, 1999). This mucoid form of the bacterium is virtually untreatable with traditional antibiotics, making it necessary to seek alternate modes of attack (Hill *et al.*, 2005). Since all pathogenic organisms need to obtain energy from host metabolites, if you cut off this energy source, the bacteria are left without the ability to produce the virulence factors contributing to their defense against the host. Thus, energy-producing metabolic pathways may provide a new drug target for treatment of *P. aeruginosa* infections.

Little is known about L-serine degradation in *P. aeruginosa.* However, genome sequence analysis reveals the presence of two L-SDs encoded by the genes *sdaA* (*PA2448)* and *sdaB* (*PA5379*) (Stover *et al.*, 2000). The *P. aeruginosa* L-SD proteins share the conserved cysteine residues involved in the stabilization of a catalytic iron sulfur center (Flint & Allen, 1996). This data indicates that these enzymes, like their prokaryotic homologs, most likely exhibit catalysis via an iron sulfur cluster. Given the similarity of the *P. aeruginosa* homologs to the L-SD proteins implicated in pathogenicity, we postulate that *P. aeruginosa* L-SD plays a similar role in the physiology of *P. aeruginosa.*

The pathogenicity of *P. aeruginosa* is of interest not only in CF patients who ultimately succumb to the infection, but also in burn patients, and in patients acquiring *Pseudomonal* nosocomial (hospital-acquired) infections. Heavy emphasis on this organism’s resistance to both bacteriostatic and bactericidal treatment has elucidated a heavy arsenal of antibiotic resistance mechanisms which make classical treatment with antibiotics ineffective at best. The failure to treat patients with these infections, fuels the drive to explore new areas upon which we might effectively hinder or even eliminate the bacterium’s growth *in vivo*. In light of the recent elucidation of the role of L-SD in the pathogenicity of several organisms (*S. pyogenes, M. bovis, C. jejuni),* we hypothesize that the L-SD in *P. aeruginosa* confers virulence. *P. aeruginosa* harbors two genes that encode L-SD, *sdaA* and *sdaB*. The aims of this study are to characterize L-serine utilization by *P. aeruginosa*, determine the contribution of *sdaA* and *sdaB* to the total L-SD activity, and investigate *P. aeruginosa* L-SD contribution to its virulence

## MATERIALS AND METHODS

### Bacterial and Nematode Strains, Plasmids and Media

Genotypes and sources of bacterial and nematode strains, plasmid genotypes and sources used in this experiment are listed in Table 1. *P. aeruginosa* and *E. coli* strains were cultured in Luria-Bertani (LB) broth or LB agar at 37 °C unless otherwise stated. Antibiotics for *E. coli* were added at the following concentrations: ampicillin (Ap), 50 µg/ml, carbenicillin (Cb), 10 µg/ml, gentamycin (Gm), 15 µg/ml, kanamycin (Km), 50 µg/ml, tetracycline (Tc), 20 µg/ml. For *P. aeruginosa* Cb, 300 µg/ml, Gm, 300 µg/ml, Nm, 150 µg/ml, and Tc, 60 µg/ml were used. Nematodes were cultured in Nematode Growth Media unless otherwise stated.

**Table 1.**
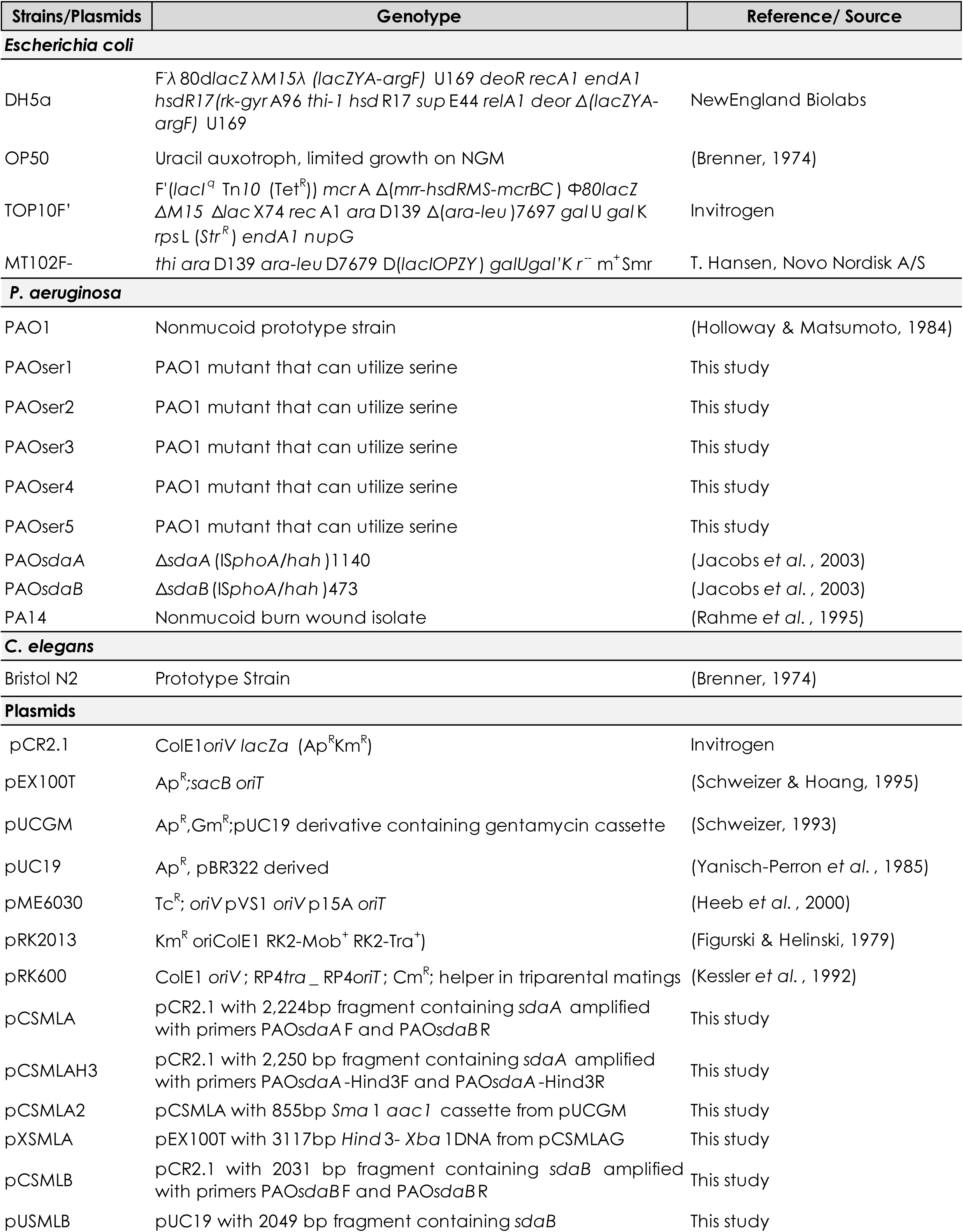
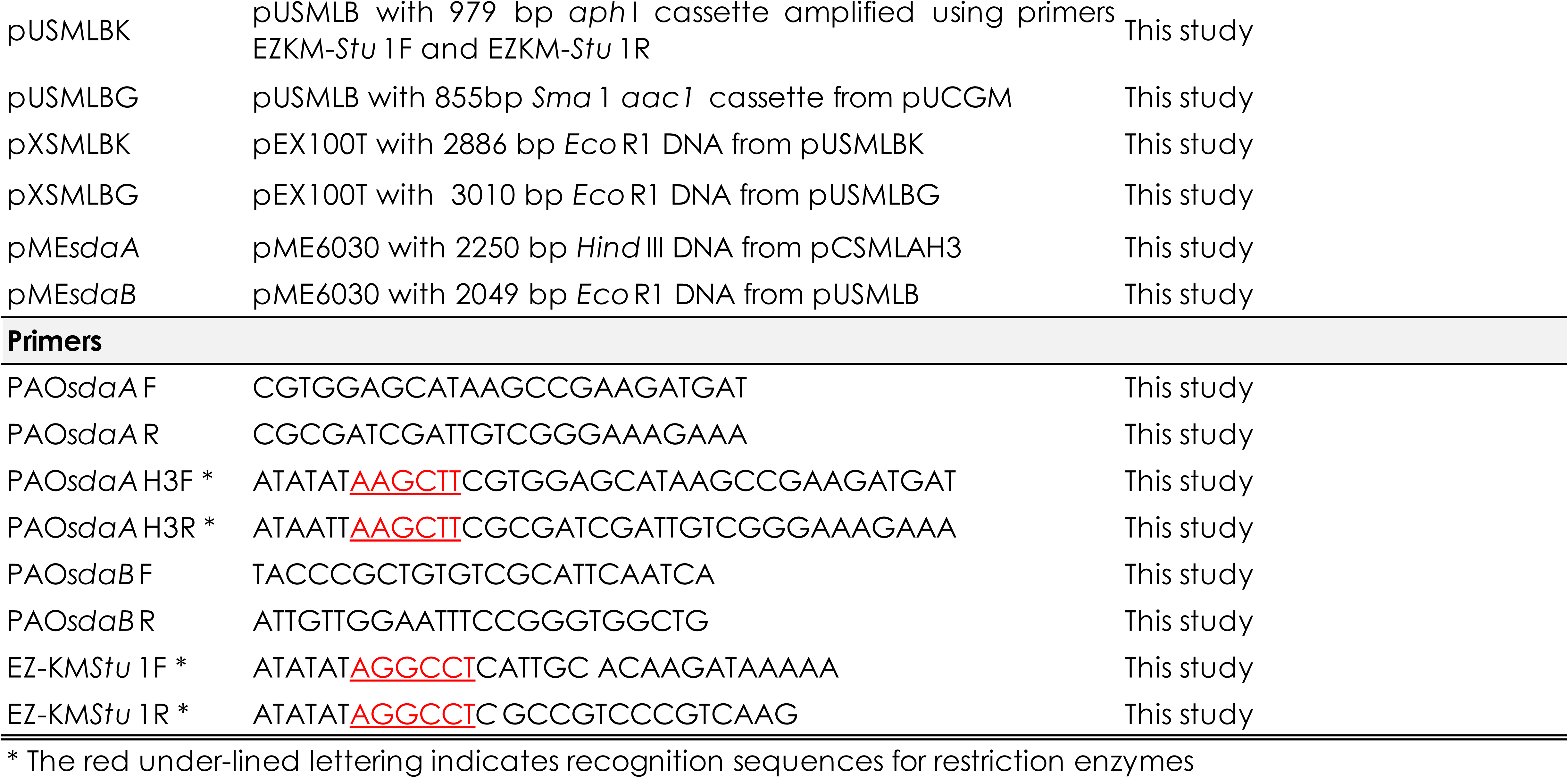
Genotypes and source of bacterial and nematode strains, plasmids, and primers

### Protein Sequence Analysis of *P. aeruginosa* L-SD

Protein sequence homology analysis was performed between the two L-SDs encoded by *P. aeruginosa* and several L-SD homologs in bacterial species of interest utilizing the program Protean, DNA STAR, 2004. These proteins and bacteria consisted of *E. coli* SdaA and SdaB, *Mycobacterium tuberculosis* SdaA, *M. bovis* SdaA, *Campylobacter jejuni* SdaA, *Streptococcus pyogenes* SdaA, and *Yersinia pestis* SdaA. Further analysis of SdaA and SdaB was performed via the computer program Protean, DNA Star, 2004. Hydrophobicity plots for SdaA and SdaB were performed utilizing the following Colorado State University website: http://www.vivo.colostate.edu/molkit/hydropathy/index.htm

### Chromosomal DNA Extraction

A modification of the Qiagen Genomic DNA Kit (Qiagen Inc., Valencia, CA) was used to extract *P. aeruginosa* chromosomal DNA (Qiagen Inc., Valencia, CA). Strains of interest were grown overnight at 37 °C. One milliliter of this culture was centrifuged and resuspended in Buffer B1 (Bacterial Lysis Buffer: 50 mM Tris-HCl, 50 mM EDTA, 0.5 % Tween-20, 0.5 % Triton X-100, pH 8.0) and 2-µl RnaseA (100 µg/ml). Twenty microliters of lysozyme (100 mg/ml) was added, followed by 45 µl Proteinase K (20 mg/ ml). The suspension was incubated at 37 °C for 30 minutes, then 175 µl of Buffer B2 (Bacterial lysis buffer: 3 M guanidine HCl, 20 % Tween-20) was added. The tubes were inverted several times and incubated at 50 °C for 30 minutes. An equal volume of phenol/chloroform was added, followed by vortexing and centrifugation. A 0.1 volume of 3 M sodium acetate and 2 volumes of ice cold 70 % (v/v) ethanol were added to the decanted supernatant and incubated for 30 minutes at −20 °C. The DNA-containing solution was vacuum dried and resuspended in 50 µl of dH_2_O.

### Gene Amplification

To obtain a sufficiently high quantity of DNA for cloning, genes of interest were amplified via the polymerase chain reaction (Techne, Albany, NY). To amplify *sdaA*, the primer pairs PAO*sdaA*F, PAO*sdaA*R and PAO*sdaA*H3F, PAO*sdaA*H3R were used (Table 1). To amplify *sdaB*, the primer pair PAO*sdaB*F and PAO*sdaB*R were used (Table 1).

The parameters used to amplify both *sdaA* and *sdaB* from *P. aeruginosa* chromosomal DNA are as follows: initial denaturation of chromosomal DNA occurred at 95 °C. This step was followed by 30 cycles of denaturation for 1 minute at 95 °C, annealing for 1 minute at 63 °C, and extension for 4 minutes at 72 °C. The final extension occurred at 72 °C for 10 minutes followed by a final hold at 4 °C.

To amplify the Km resistance cassette *ahc*I from the EZ-Km transposon kit the same PCR parameters were used except for the primer extension time that was set at 1 minute. The primers used were EZ-Km*Stu*IF and EZ-Km*Stu*IR (Table 1).

### DNA Isolation

In order to obtain pure DNA, the fragments of interest were separated via electrophoresis on a 1 % agarose gel made with TBE buffer. DNA was stained with either ethidium bromide or the non-carcinogenic alternative SYBR SAFE (Invitrogen, Carlsbad, CA) and visualized under UV light. DNA fragments were excised from the gel with a scalpel. The gel fragment containing the DNA was weighed out and three volumes of Buffer QG (Solubilization and Binding Buffer) were added to the gel fragment. The mix was incubated at 50 °C until the solution was homogeneous; one gel volume of isopropanol was added and the mixture was vortexed. This solution was applied to the QIAspin column filter (Qiagen Inc., Valencia, CA) and centrifuged for 1 minute at 13,000 rpm. The filtrate was discarded and the column was washed using 750 µl of Buffer PE (100 ml Buffer PE, 400 ml ethanol) and centrifuged for one minute. The filtrate was discarded and a second spin performed to remove residual ethanol in the column. Thirty to 50 µl of Buffer EB (Elution Buffer; 10 mM Tris-HCl, pH 8.5) was added. Following a 1-minute incubation period, a final spin eluted the purified DNA of interest.

### Plasmid DNA Extraction

In order to extract plasmids of interest from the cells harboring them, the Qiaprep Spin Miniprep Kit was used (QIAGEN Inc.,Valencia, CA) Strains harboring the plasmids were grown overnight in 10 ml LB with appropriate antibiotic. The cells were harvested by centrifugation at 8,000 x g for 6 minutes. The pellet was resuspended in 500 µl Buffer P1 (Resuspension buffer; 50 mM Tris-Cl, pH 8.0, 10 mM EDTA; 100 µg/ml RNaseA) followed by the addition of 500 µl of Buffer P2 (Lysis Buffer; 200 mM NaOH, 1 % Sodium Dodecyl Sulfate). The tubes were inverted to gently mix the solution and incubated at room temperature (RT) for 5 minutes before adding 750 µl of Buffer N3 (Neutralization Buffer; 3.0 M potassium acetate, pH 5.5) to neutralize the reaction. Two ml aliquots were transferred to micro centrifuge tubes and then centrifuged at 13,000 x g. The supernatant containing the DNA was decanted into a QIA prep spin column filter. The column was centrifuged and washed with 750 µl of Buffer PE (100 ml Buffer PE, 400 ml 100 % ethanol). The DNA was eluted into a new micro centrifuge tube after incubation for one minute at 25 °C with 50 µl of Buffer EB (Elution Buffer; 10 mM Tris-HCl, pH 8.5) and a final spin down.

### Preparation of Competent Cells and Chemical Transformation

To make chemically competent *E. coli* DH5α cells for plasmid transformation, DH5α was cultured in 5 ml of LB broth at 37 °C with shaking. After overnight incubation a 1:100 subculture into 50 ml of LB broth was incubated till an OD_600_ value of 0.5 to 1.0 was reached. The cells were centrifuged at 6000 rpm in a refrigerated micro centrifuge set to 0 °C. The supernatant was decanted and to the cells were added 25 ml of ice-cold 0.1 M CaCl_2_. The suspension was mixed via tube inversion. The cells were then chilled on ice for 15 minutes and centrifuged once more as stated above. The cells were then resuspended in 3.3 ml of sterile, ice-cold 0.1M CaCl2-15% glycerol, and left on ice for 4 to 20 hours. Fifty microliter aliquots were transferred to micro centrifuge tubes and either used immediately to transform plasmid DNA of interest or stored at −70 °C up to several months.

Ten µl of ligated or vector DNA was added to the competent cells and incubated on ice for 15 minutes. The cells were then heat shocked for 2-3 minutes at 37 °C and immediately placed on ice for 5 minutes. One ml of LB broth was added and the culture placed in a shaking incubator at 37 °C for one hour to allow time for transformant recovery to take place. Post incubation, 100 µl of this mix was plated on selective media. The remaining cells were spun down, resuspended in 100 ul of LB, and also plated on selective media. The cells were then incubated overnight at 37 ° C and observed the following day.

### Preparation of Electrocompetent Cells and Electroporation

In order to prepare electrocompetent cells 500 µl of overnight culture was inoculated into 500 ml of LB broth. The cells were grown at 37 °C with shaking at 250 rpm to an OD_600_ of approximately 0.6. The cells were chilled for 20 minutes; then centrifuged at 4000 g for 15 minutes at 4 °C. The cell pellet was resuspended and washed with 500 ml of ice-cold 10 % (v/v) glycerol, centrifuged and resuspended in 20 ml of the same medium. This suspension was centrifuged and resuspended in 2 ml ice cold 10 % glycerol. Forty microliter aliquots were transferred to micro centrifuge tubes for storage at −70 °C for later use.

Frozen electrocompetent cells were thawed (40 µl) and 2 µl of the DNA of interest was added. The cells were incubated for 1 minute on ice and transferred to a 0.2 cm gene pulser cuvette (BIORAD, Hercules, CA), at which point the tube was placed in the electroporation chamber and electroporated at setting *E. coli* 2 on the BIORAD Micropulser (BIORAD, Hercules, CA). Immediately after pulsing, 1 ml SOC medium (2.0 % tryptone, 0.5 % yeast extract, 10 mM MgCl_2_, 10 mM NaCl, 2.5 mM KCl, 10 mM MgSO_4_, 20 mM glucose) was added and the suspension transferred to a new micro centrifuge tube. This culture was incubated for 30 min at 37 °C with shaking. It was spun down and resuspended in 100 µl of LB medium. Fifty microliters of this culture was spread on selective media and incubated overnight at 37 °C.

### Plasmid Construction

Plasmid constructs were made which allow for selective inactivation of *sdaA* and *sdaB* within the *P. aeruginosa* chromosome.

### Construction of *sdaA*::*aacCI*, pXSMLA

To force homologous double crossover recombination at the wild type *sdaA* locus in the *P. aeruginosa* chromosome, a suicide vector (capable of chromosomal allelic replacement) carrying an insertionally inactivated Δ*sdaA:aac1* cassette was constructed.

The first step in suicide vector formation involved amplification of the *sdaA* gene with primers PAO*sdaA*F and PAO*sdaA*R (Table 1). Taq DNA polymerase leaves 3’deoxyadenosine residues during the amplification reaction and pCR2.1 has engineered 3’ deoxythymidine residues allowing for easy annealing and ligation using topoisomerases to occur (Invitrogen, Carlsbad, CA.). The 2244 bp amplicon was cloned into the pCR2.1 vector (Invitrogen, Carlsbad, CA) (Figure 3). The reaction involved a 5-minute incubation at 25 °C with 4 µl of PCR-amplified DNA, 1 µl of 10X Reaction Buffer, and 1 µl of the vector pCR2.1. The ligation mix was used to transform TOP10F’ cells (Invitrogen, Carlsbad, CA). Upon DNA insertion into pCR2.1’s cloning site, the l*acZ* gene was disrupted, allowing selection of recombinants based on blue-white screening. Transformants were plated on LB containing Ap (50 µg/ml) plates that were spread with 40 µL of 5-Bromo-4-chloro-3-indolyl-β-D galactopyranoside (X-gal, 40 mg/ml) and 40 µl Isopropyl β-D-thiogalactopyranoside (IPTG, 100 mM; only white transformant colonies were selected. Confirmation of the correct clone was carried out via gel electrophoresis of the *Eco*R1 digest on a 1 % agarose gel in TBE Buffer. The cloned pCR2.1 derivative with the *sdaA* gene insert was named pCSMLA (Figure 3).

**Figure 3.**
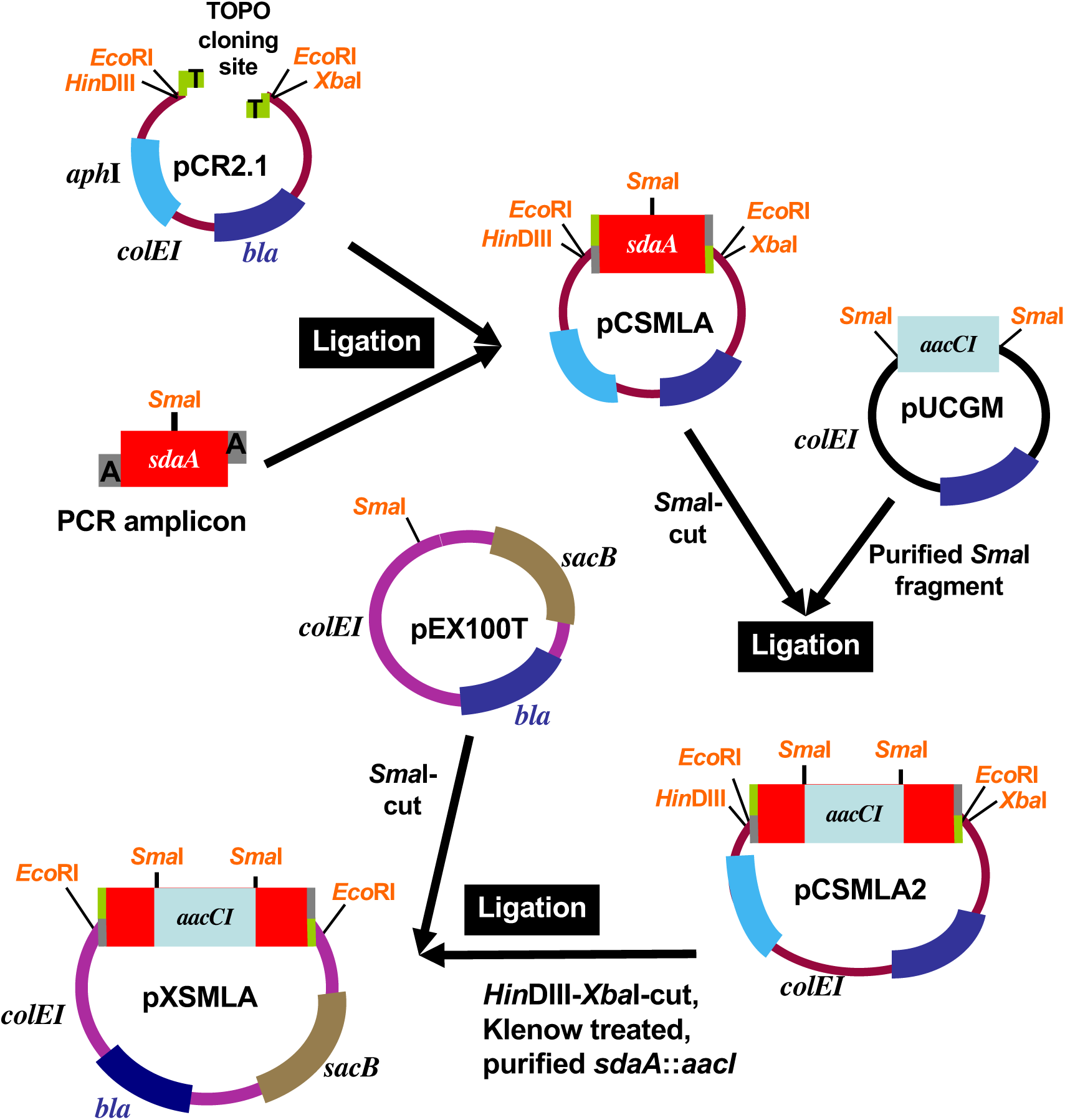
Construction of a suicide vector containing insertional inactivated *sdaA* with *aacCI.* The *sdaA* gene was PCR amplified from the PAO1 chromosome and annealed to pCR2.1 forming pCSMLA. The *Sma*I fragment from pUCGm containing *aac*I was cloned into a unique *Sma*I site on the *sdaA* gene on pCSMLA forming pCSMLA2. A 3,117-bp *Hin*dIII-*Xba*I-Klenow treated fragment containing the *sdaA*::*aacC1* fragment was sub cloned into a *Sma*1 site on the suicide vector pEX100T that contains *sacB* gene for counter selection and named pXSMLA.

This plasmid was derived via sub cloning the 855-bp *Sma*I fragment containing *aac*I encoding gentamycin resistance, from pUCGM (Schweizer, 1993) into a unique *Sma*1 site on the *sdaA* gene on pCSMLA (Figure 3). The blunt end ligation was carried out via overnight incubation with 10 µl Ligase Buffer, 2 µl T4 ligase, and 0.01 µM of *Sma*1 digested *aac*I and 0.090 µM of the *aac*I cassette, 10:1 molar ratio (New England Bio labs Beverly, MA). Electrocompetent DH5α cells were electroporated with 2 µL of the ligation mix, and transformants selected on LB plates containing Ap.

Transformants were grown in LB Broth with Gm (15µg/ml). Confirmation of the correct clone was carried out via gel electrophoresis of the *Eco*R1 digest on a 1 % agarose gel in TBE Buffer. The pCSMLA derivative with *sdaA:aac*I was named pCSMLA2 (Figure 3).

A 3,117-bp *Hin*dIII-*Xba*I-Klenow treated fragment containing the *sdaA:aac*I fragment was sub cloned into a *Sma*I site on pEX100T (Figure 3). The transformants were selected on LB agar plates containing Ap (50 µg/ml). Confirmation of the correct clone was carried out via gel electrophoresis of the *Fsp*I digest on a 1 % agarose gel in TBE Buffer. The pEX100T derivative was named pXSMLA.

### Construction of *sdaB*::*aphI*, pXSMLBK

To force homologous double crossover recombination at the wild type *sdaB* loci in the *P. aeruginosa* chromosome, a suicide vector carrying an insertionally inactivated Δ*sdaB:aph1* cassette, pXSMLBK, was constructed (Figure 4). The *sdaB* gene was PCR amplified from the PAO1 chromosome using the primer pair PAO*sdaB*F and PAO*sdaB*R (Figure 4, Table 1). The amplicon was sub cloned into pCR2.1 forming pCSMLB (Figure 4).

**Figure 4.**
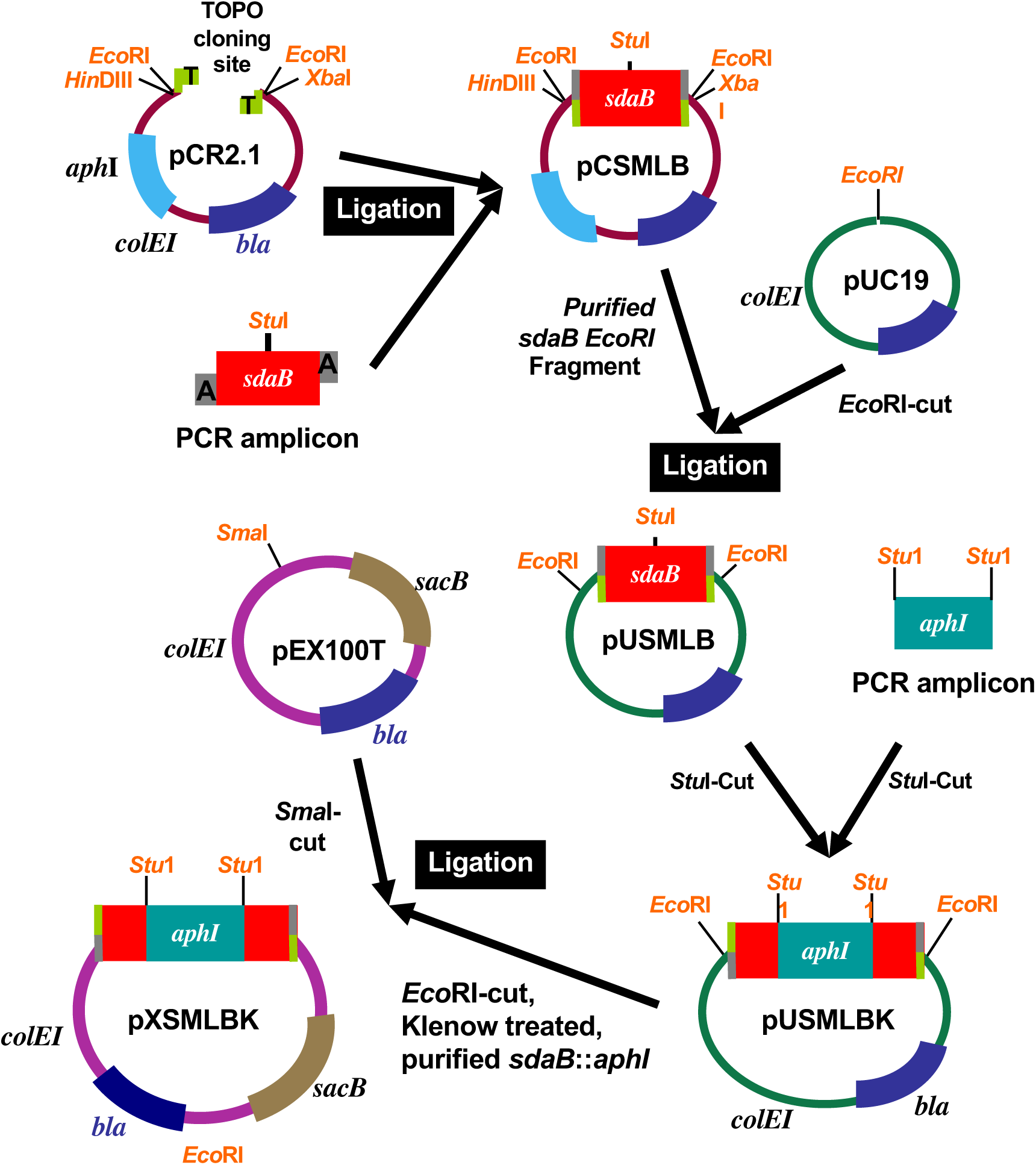
Construction of a suicide vector containing insertional inactivated *sdaB* with *aphI.* The *sdaB* gene was PCR amplified from the PAO1 chromosome and annealed to the vector pCR2.1 forming pCSMLB. An *Eco*R1 fragment containing the *sdaB* gene from pCSMLB was sub cloned into at the *Eco*RI site on pUC19 forming pUSMLB. The 979-bp *Stu*I digested *aphI* PCR amplicon from EZ-KM transposon was ligated to *Stu*I digested pUSMLB forming pUSMLBK. The 3010-bp *Eco*R1-Klenow treated *sdaB*::*aph*I from pUSMLBK was sub cloned into the *Sma*1 site of the suicide vector pEX100T that contains *sacB* gene for counter selection forming pXSMLBK.

The 2049-bp *Eco*R1 DNA bearing *sdaB* from pCSMLB was sub cloned into *Eco*RI-cut pUC19 forming pUSMLB (Figure 4). The switch from pCR2.1 to pUC19 was carried out because pUC19 lacks the kanamycin resistance marker (*aph*I). A second *aph*I cassette was later used to insertionally inactivate the *sdaB* gene.

A 979-bp fragment containing *aph*I encoding kanamycin resistance was amplified from the EZ-transposon kit using the primer pair EZ-Km*Stu*IF and EZ-Km*Stu*IR (Figure 4, Table 1). The amplicon was digested with *Stu*1 and sub cloned into a *Stu*I cut pUSMLB forming pUSMLBK (Figure 4). A 3010-bp *Eco*R1-Klenow fragment containing *sdaB:aph*I from pUSMLBK was ligated to a *Sma*I cut pEX100T forming pXSMLBk. (Figure 4). Transformants were selected on LB + Km (50 µg/ml) and the clone confirmed via *Fsp*I digest.

### Construction of *sdaB*::*aacCI*, pXSMLBG

Another suicide vector, pXSMLBG, with *sdaB* insertionally inactivated using a gentamycin cassette (*aac*I) was constructed (Figure 5). A 855-bp *aac*I DNA fragment was sub cloned into the *Stu*1 site of pUSMLB forming **pUSMLBG** (Figure 5). This plasmid was then digested with *Eco*R1, releasing two fragments of 2686 bp and 2886 bp respectively. The 2886-bp fragment containing the *sdaB*:*aac*I cassette was ligated to *Sma*1 cut pEX100T forming **pXSMLBG** (Figure 5). The presence of the right clone was confirmed via *Fsp*I digestion.

**Figure 5.**
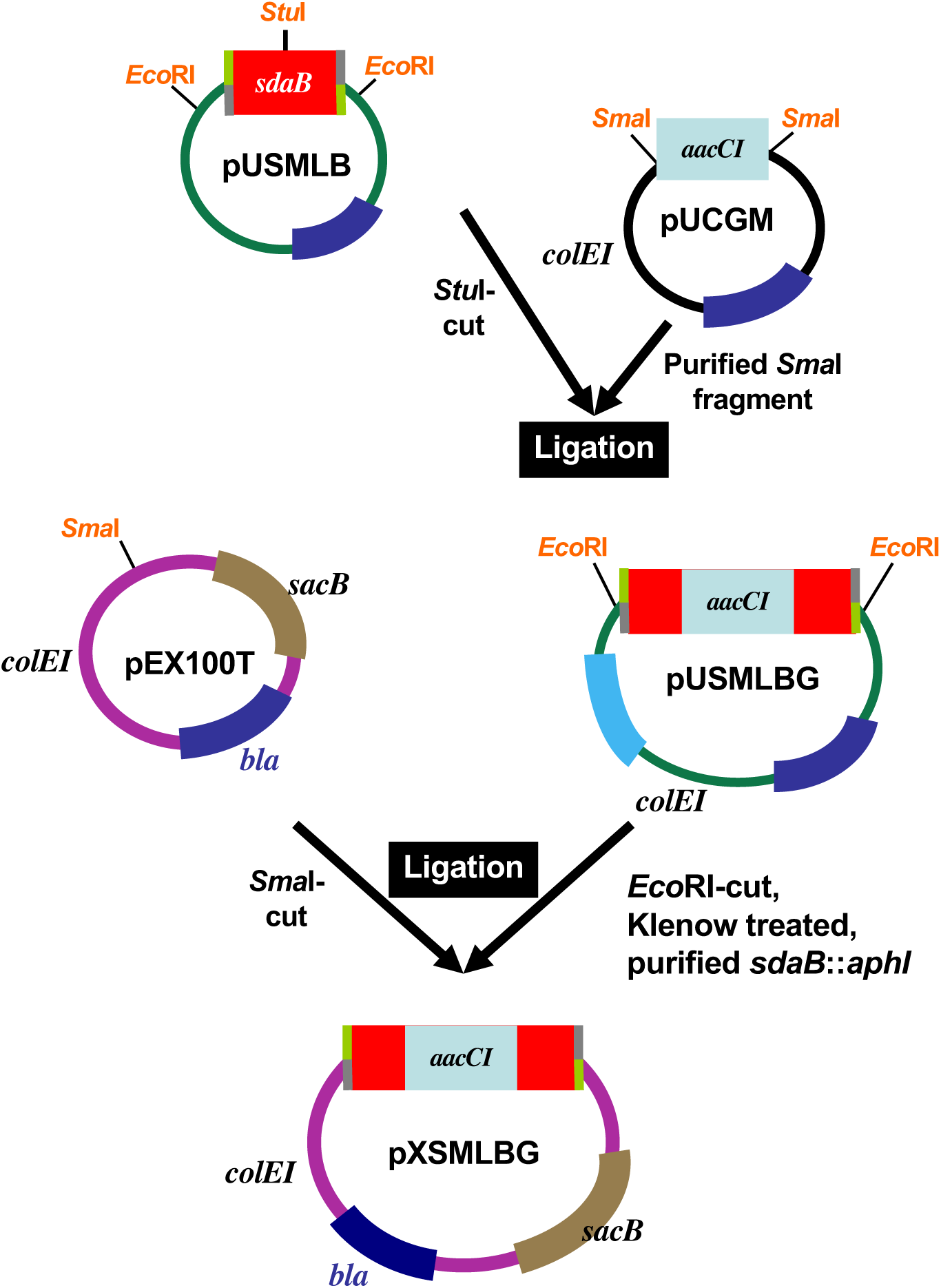
Construction of a suicide vector containing insertional inactivated *sdaB* with *aacCI.* The 855-bp *Sma*I fragment containing *aacCI* cassette from pUCGm was ligated to *Stu*1 digested pUSMLB (see Figure 6) forming pUSMLBG. The *Eco*R1-Klenow treated *sdaB*::*aacCI* from pUSMLBG was sub cloned into the *Sma*1 site of pEX100T forming pXSMLBG.

### Construction of Complementing Plasmids

In order to construct complementing plasmids with *sdaA* and *sdaB*, the stable, broad host range, low copy plasmid, pME6030, was used (Heeb *et al.*, 2000).

### pME*sdaA*

A 2250-bp *Hind*III fragment bearing *sdaA* was purified from pCSMLAH3 and ligated to *Hin*dIII-cut pME6030 (Figure 6). Confirmation of the correct clone was carried out via gel electrophoresis of the *Hind*III digest on a 1 % agarose gel in TBE Buffer. The pME6030 derivative with *sdaA* was named pME*sdaA* (Figure 6).

**Figure 6.**
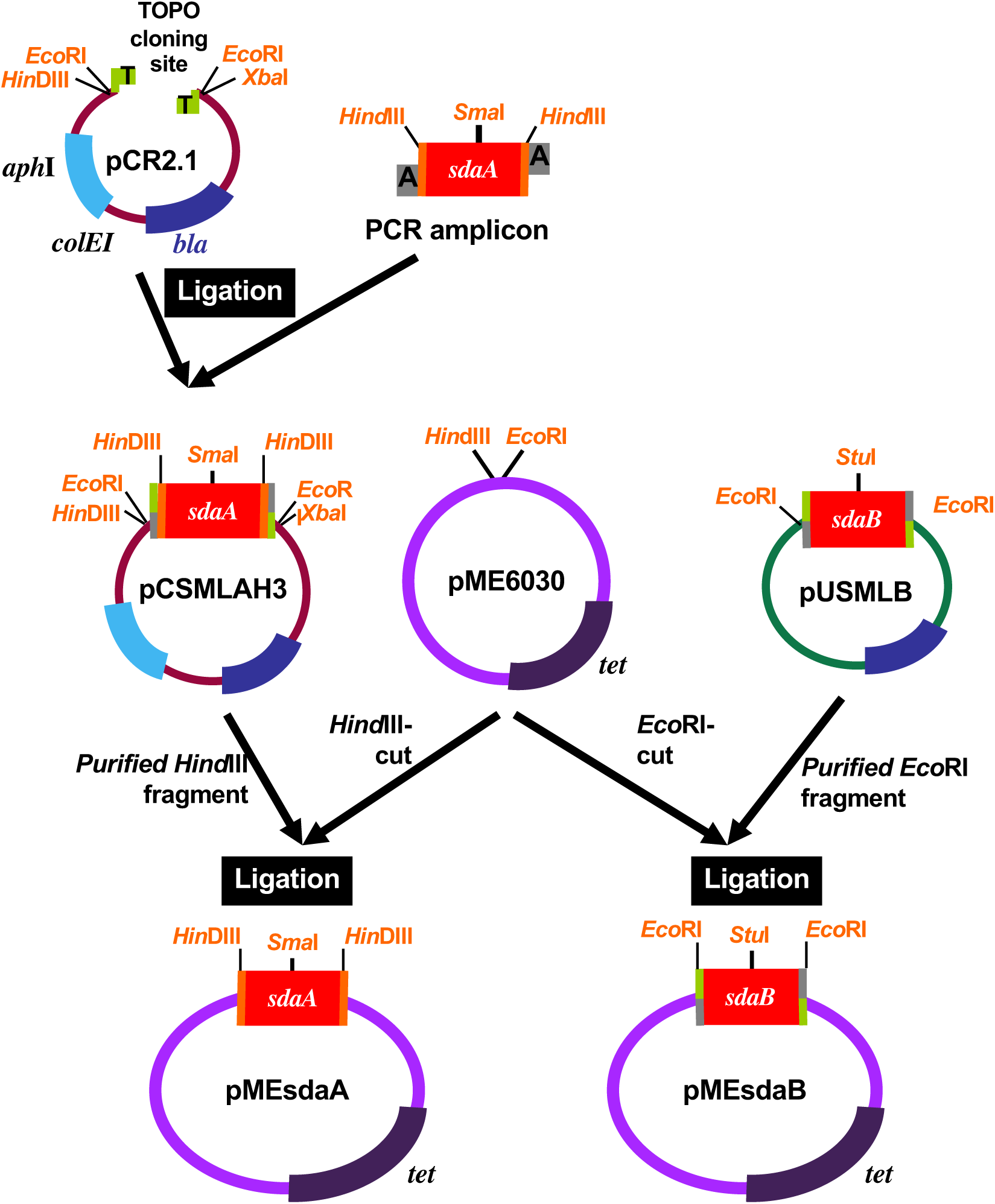
Construction of complementing plasmids. In order to obtain plasmids harboring *sdaA* and *sdaB* that are efficiently maintained for phenotypic complementation in *P. aeruginosa*, the broad host-range plasmid pME6030 was used. *Hind*III -cut pME6030 was ligated a 2250-bp *Hin*dIII fragment harboring *sdaA* from pCSMLAH3 forming pME*sdaA*. *Eco*RI-cut pME6030 was ligated to a 2049-bp *Eco*R1 fragment harboring *sdaB* from pUSMLB (Figure 6) generating pME*sdaB.*

### pME*sdaB*

A 2049-bp *Eco*R1 fragment harboring *sdaB* was transferred from pUSMLB to *Eco*RI-cut pME6030 (Figure 6). Confirmation of the correct clone was carried out via gel electrophoresis of the *Pvu*II digest on a 1 % agarose gel in TBE Buffer. The pME6030 derivative with *sdaB* was named pME*sdaB* (Figure 6).

### Construction of Suicide Vectors for Mutant Construction

#### Sub cloning *sdaA* and *sdaB* from the PAO1 Chromosome

The *sdaA* gene was successfully amplified from the PAO1 chromosome using two different primer pairs. The 2346-bp fragment amplified using PAO*sdaA*H3F and PAO*sdaA*H3R was referred to as *sdaAH*3 (Figure 7A, Lane 2). A second fragment containing *sdaA* was amplified (2334 bp) with the primer pairs, PAO*sdaA*F and PAO*sdaA*R (data not shown). The *sdaB* gene was also successfully amplified (2049 bp) from the PAO1 chromosome using the primers PAO*sdaB*F and PAO*sdaB*R (Figure 8, Lane 2).

**Figure 7.**
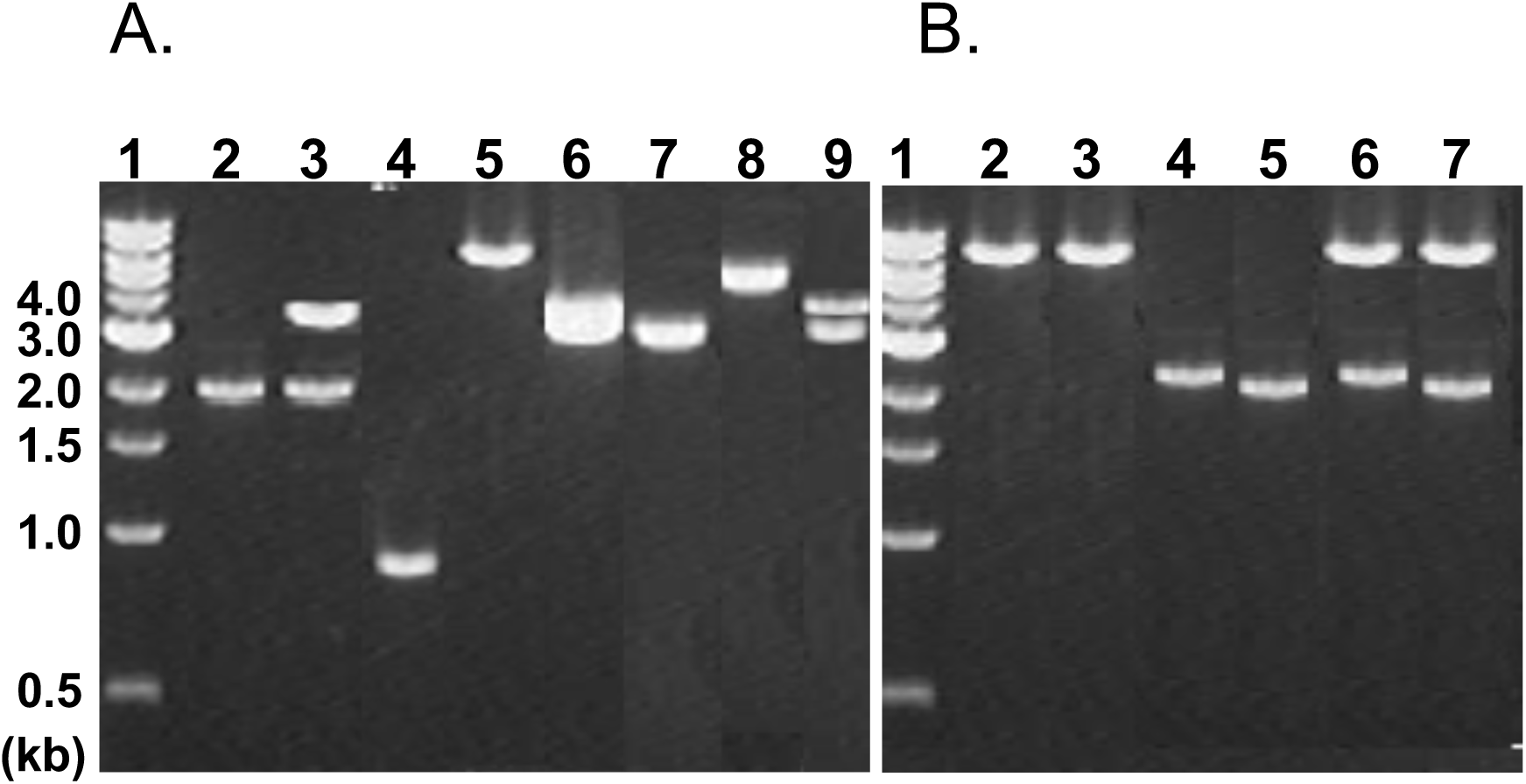
Verification of *sdaA* plasmids and complementing plasmids constructed using 1.0 % agarose electrophoresis. **Lanes 1, DNA** Ladder N3232S (New England Bio labs); **Panel A) 2.** *sdaA* gene amplified from the PAO1 chromosome using the primers PAO*sdaA*H3F and PAO*sdaA*H3R (2346bp); **3.** *Hin*dIII-cut pCSMLAH3 yielding three fragments of 67, 2250, and 3882 bp; 4. 855-bp *aac*I from pUCGm; **5.** *Sma*I-cut pCSMLA (6199 bp); **6.** *Hind* III-*Xba*I-cut pCSMLA2 yielding two 3211 and 3819 bp fragments; 7. 3211 bp *sdaA:aac*I from pCSMLA2; **8.** *Sma*I-cut pEX100T (5846 bp); **9.** *Fsp*I-cut pXSMLA yielding two confirmatory linear DNA 4183 bp and 4874 bp fragments. **Panel B) 2.** 8021-bp *Eco*RI-cut pME6030; **3.** 8021-bp *Hin*dIII-cut pME6030; **4.** 2049-bp *Eco*RI fragment with *sdaB* from pUSMLB; **5.** 2250-bp *Hin*dIII fragment with *sdaA* from pCSMLAH3; **6.** *Eco*RI-cut pME*sdaB* yielding two fragments of 8021 bp and 2049 bp; **7,** *Hin*dIII-cut pME*sdaA* yielding two fragments of 8021 bp and 2250 bp.

**Figure 8.**
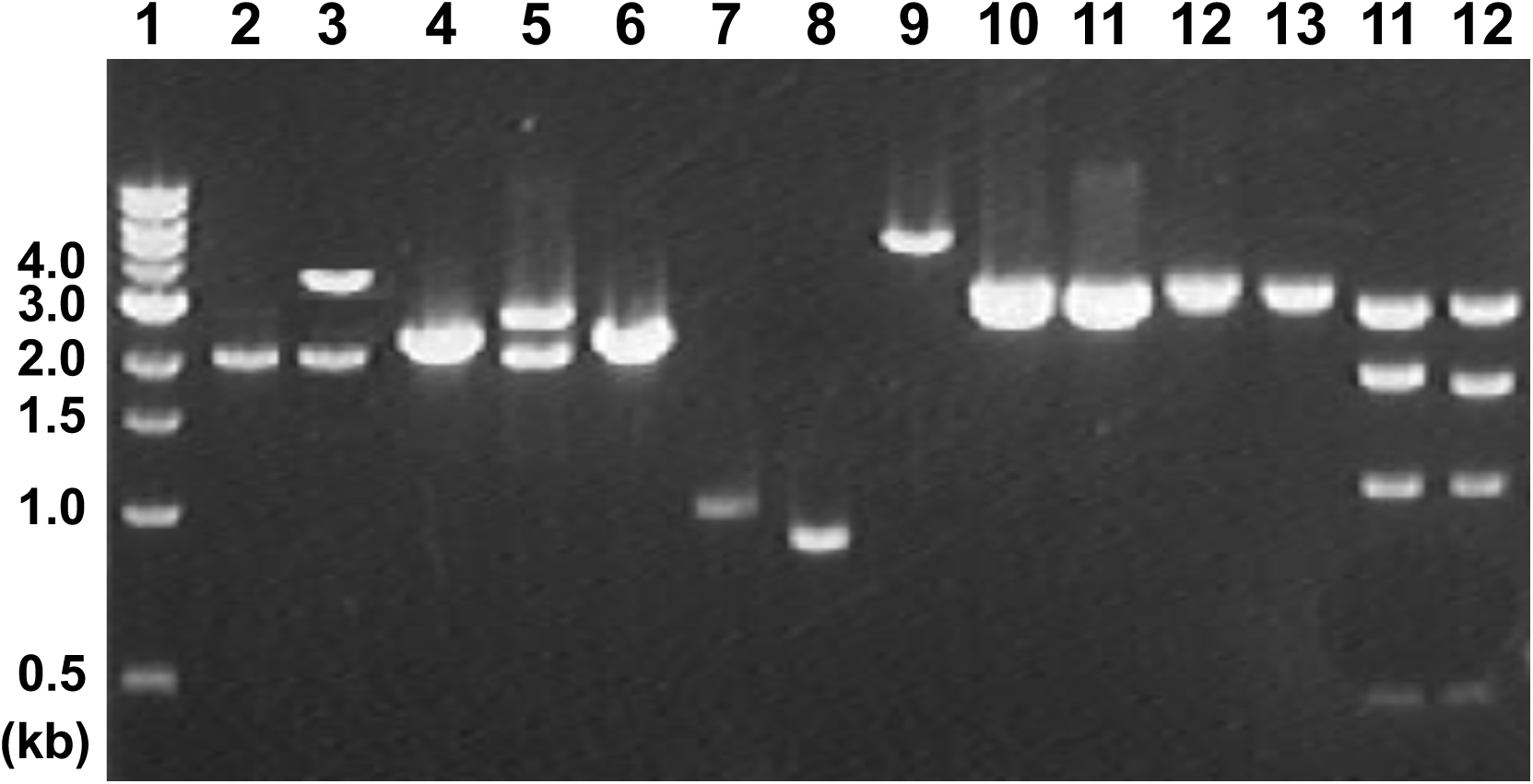
Verification of *sdaB* plasmids constructed using 1.0 % agarose electrophoresis. Lanes 1. DNA Ladder N3232S (New England Bio labs); **2.** *sdaB* gene amplified from the PAO1 chromosome using the primers PAO*sdaB*F and PAO*sdaB*R (2043 bp); **3.** *Eco*RI-cut pCSMLB yielding two fragments of 2049 and 3913 bp; **4.** 2049-bp *Eco*RI from pCSMLB with *sdaB;* **5.** *Eco*R1-cut pUSMLB, yielding the expected 2049 and 2686 bp fragments; **6.** 2049-bp *Eco*RI fragment harboring *sdaB* from pUSMLB; **7.** 979-bp amplicon from EZ-transposon kit harboring *aphI*; **8.** 855-bp *aacCI* amplicon from pUCGm, **9.** *StuI*-cut pUSMLB (4735 bp); **10.** *Eco*RI-cut pUSMLBK yielding two fragments of 2686 bp and 3010 bp; **11.** *Eco*RI-cut pUSMLBG yielding two fragments of 2686 and 2904 bp; **12.** 3010-bp fragment from pUSMLBK with *sdaB*::*aphI*; **13.** *sdaB::aacCI* from pUSMLBG; **14.** *FspI*-cut pXSMLBK yielding four fragments of 363, 1023, 1714, and 2596 bp; and **15.** *FspI*-cut pXSMLBG yielding four fragments of 363, 1023, 1608, and 2596 bp.

All three amplicons (*sdaA*, *sdaA*H3, and *sdaB*) were ligated to vector pCR2.1 (forming pCSMLA, pCSMLAH3, and pCSMLB) (Figure 3, 4, 5). *Eco*R1 restriction enzyme digest of pCSMLA yielded three confirmatory fragments of 743, 1515, and 3913 bp (data not shown). A *Hin*dIII restriction enzyme digest of pCSMLAH3 yielded three confirmatory fragments of 67, 2250, and 3882 bp (Figure 7A, Lane 3). Finally, *Eco*R1 digest of pCSMLB yielded two confirmatory fragments of 2049 bp and 3913 bp (Figure 8, Lane 3).

#### Confirmation of Complementary Plasmids

The 2250-bp *Hin*dIII fragment containing *sdaA* from pCSMLAH3 was ligated *Hin*dIII-cut pME6030 (Figure 7B, Lane 2,4) (Figure 3). *Hin*dIII digestion of the clone (pME*sdaA*), confirmed the presence of two amplicons 2250 and 8021 bp (Figure 7B, Lane 6). The *sdaB* amplicon was cloned into the vector pCR2.1 yielding pCSMLB (Figure 4). pCSMLB yielded two fragments of 2049 and 3913 bp when digested with *Eco*RI (Figure 8, Lane 3). The 2049-bp *Eco*R1 DNA was successfully ligated to pME6030-*Eco*R1 (Figure 7B, Lane 3) (Figure 6). pME6030-*sdaB* (pME*sdaB*) transformants were characterized by Tc^R^ and screened via an *Eco*R1 digest revealing two bands of 2049 bp and 8021 bp (Figure 7B, Lane 7).

#### Transfer of *sdaB* to a Km^S^ Plasmid

The 2049-bp *Eco*R1 fragment from pCSMLB (Figure 4) was successfully ligated to *Eco*RI-cut pUC19. The transformant phenotype for pUC19-*sdaB* (pUSMLB) was the following: Ap^R^, Km^S^, and white colony formation on growth with X-gal. *Eco*R1 digest of pUSMLB (Figure 4), yielded the expected fragments of 2049 bp and 2686 bp (Figure 8, Lane 5).

#### Insertional Inactivation of *sdaA* and *sdaB* via an *aacCI* Cassette

The 855-bp *Sma*I fragment from pUCGm containing the *aacCI* gene encoding gentamycin resistance, was ligated to *Sma*1-cut pCSMLA (Figure 3) and a *Stu*I cut pUSMLB (Figure 5) (Figure 8, Lane 7 and 9, Figure 7A, Lane 5). *Hind*III*-Xba*I double digest of the Gm^R^ pCSMLA::*aacC*I plasmid (pCSMLA2), yielded two confirmatory fragments of 3211-bp and 3819-bp (Figure 7A, Lane 6). pUSMLB:*aac*I (pUSMLBG) yielded two confirmatory fragments of 2686 and 2904 bp when cut with *Eco*RI (Figure 8, Lane 11).

#### Insertional Inactivation of *sdaB* via an *aphI* Cassette

The 979-bp *aph*I gene, encoding kanamycin resistance, was successfully amplified from the EZ transposon kit using the primer pair EZKm*Stu*IF and EZKm*Stu*1R (Figure 8, Lane 7). The *aphI* amplicon was restricted with *Stu*I and ligated to *Stu*I cut pUSMLB yielding transformants with the following phenotype: Ap^R^, Km^R^, white on X-gal selection (Figure 8, Lane 9). *Eco*R1 digest of the pUSMLB-*aph*I clone (pUSMLBK) revealed two confirmatory fragments of 2686 bp and 3010 bp (Figure 4) (Figure 8, Lane 10).

#### Suicide Vectors Bearing *sdaA*::*aacCI, sdaB*::*aacCI,* and *sdaB*::*aphI*

Fragments containing *sdaA*::*aacCI, sdaB*::*aacCI, and sdaB*::*aphI* were sub cloned into the suicide vector pEX100T from pUSMLBK, pUSMLBG, and, pCSMLA2 respectively (Figure 8, Lane 12 and 13, Figure 7A, Lane 7). As expected, a *Hind*III and *Xba*I double digested pCSMLA2 yielded two linear DNA bands of 3211 and 3819 bp (Figure 7A, Lane 6). *Eco*RI digestion of pUSMLBG, yielded two linear DNA fragments of 2686 bp and 3010 bp (Figure 8, Lane13). Similarly, *Eco*R*1* digested of pUSMLBK yielded two linear DNA bands of 2686 bp and 2904 bp (Figure 8, Lane 12).

The blunt-ended fragments representing the *sdaA*::*aacCI*, *sdaB*::*aacCI* and the *sdaB*::*aphI* cassettes of 3211 bp, 2904 bp, and 3010 bp, respectively, were ligated to *Sma*I-pEX100T (Figure 8, Lane 12 and 13, Figure 7A, Lane 7). Transformants were Ap^R^, Gm^R^, and appeared white on X-gal selection. *F*s*p*I digestion of the pEX100T-*sdaA::aacCI* clone (pXSMLA) (Figure 3) yielded two confirmatory linear DNA fragments of 4183 bp and 4874 bp (Figure 7A, Lane 9). *Fsp*I digest of the pEX100T-*sdaB*::*aacCI* clone (pXSMLBG) (Figure 5) yielded four confirmatory linear DNA fragments of 363, 1023, 1608, and 2596 bp when subjected to gel electrophoresis (Figure 8, Lane 15). *Fsp*I digest of the pEX100T-*sdaB*::*aphI* clone (pXSMLBK) (Figure 4) yielded four confirmatory linear DNA fragments of 363, 1023, 1714, and 2596 bp.

#### Determination of the Ability of *P. aeruginosa* to Grow on L-Serine

In order to determine if *P. aeruginosa* could utilize serine as the sole carbon source, all strains of interest (PAO1, PAO*sdaA*,and PAO*sdaB*) were grown overnight in NIV-Glucose (NIVGlu) media broth. Strains PAO1, PAO*sdaA*, and PAO*sdaB* were grown overnight in LB media and sub cultured in NIV glucose media overnight. Cells were resuspended in saline, diluted 10^-6^, and plated on differential media which included NIV Glu, NIV SGL (Serine, Glycine, and Leucine), NIV SG (Serine and Glycine), NIV SL (Serine and Leucine), NIV S (Serine), NIV G (Glycine), NIV L (Leucine). As a control the cells were also plated on NIV and NMM media. Colonies were observed for three days and colony size and count recorded three days’ post incubation.

#### Serine Efficient Mutant Strain Isolation and Characterization

Mutant strains capable of growth on minimal media with L-serine as the sole carbon source were isolated via prolonged growth and continuous restreaking on NIV S media. The purified mutant strains were grown for 36 hours in NIV S media at which point in time the OD_600_ value was recorded. These mutants were then subjected to L-SD assays after 14 hr. growth on NIV S and NIV SGL media.

#### *P. aeruginosa* Growth Curves in Minimal Media

PAO1, PAO*sdaA,* and PAO*sdaB* were streaked on LB agar plates +/- the addition of antibiotics, and incubated overnight. Single colonies were cultured in NIV Glu overnight. The cells were spun down at 5000 rpm for 6 minutes at 4 °C. The supernatant was decanted and the cells washed twice with sterile dH_2_O. The cells were then resuspended in the appropriate minimal media. The cultures were incubated at 37 °C with aeration. OD_600_ values were recorded every hour for 16 hours.

#### L-SD Assay Optimization

To establish a method of analyzing the amount of L-serine being utilized within a *P. aeruginosa* whole cell, the *E. coli* L-SD assay was modified (Newman & Kapoor, 1980; Isenberg & Newman, 1974). PAO1 was grown in NIV GluG and NIV SGL media for 12 hours. The L-SD assay described for *E. coli* was carried out with the reaction being stopped at 10 minute intervals for 100 minutes (Isenberg & Newman, 1974).

#### *P. aeruginosa* L-SD Assay

All strains of interest were grown in LB media with appropriate antibiotics overnight at 37 °C. The next day a 1/100^th^ subculture was transferred to NIV media (Serine, Glycine, and Leucine (SGL), Serine and Glycine (SG), Serine (S), Glucose and Glycine (GluG), Glucose and Leucine (GluL), Glucose (Glu), and LB Media). At 14 hr, the culture was cooled in an ice water suspension for one minute with swirling. The culture was centrifuged and resuspended in KPO_4_ buffer (50 mM, pH 7.5). The OD_600_ value of the suspension was adjusted to 0.300 +/- 0.010. Three hundred microliters of the adjusted cell suspension were added to five pre-warmed test tubes (1 hr. at 37 °C) containing 180 µl of 50 mM KPO_4_ with or without the addition of 20 µl of 10 % L-serine. Twenty microliters of toluene were added and the test tubes were briefly vortexed and placed in a 37 °C incubator without aeration for 50 minutes. At 50 minutes, 900 µl of dinitrophenylhydrazine (DNPH) solution was added, and the test tubes were incubated at 25 °C for an additional 20 minutes. At twenty minutes the reaction was stopped via addition of 1.9 ml of NaOH (10 % w/w). The OD_540_ values were read on the BIORAD SmartSpec3000 and converted to µg of pyruvate via the pyruvate standard curve described below.

During the assay L-serine is converted to pyruvate by *P. aeruginosa* during the 50 minute incubation at 37 °C. DNPH is allowed to react with the pyruvate in solution. and upon NaOH addition, a colorimetric change is observed. The extent of the change correlates with the amount of pyruvate produced and hence the amount of L-serine deaminated by the *P. aeruginosa* cells.

#### Pyruvate Standard Curve

Varying concentrations of sodium pyruvate (0, 0.5, 1, 2, 3, 4, and 5 µg/ml) were assayed to create a standard curve by which L-SD assay OD_540_ values could be converted to µg of pyruvate in solution. To do this, 900 µL of DNPH was added to 1 mL of the above stated sodium pyruvate solutions. The mixture was allowed to incubate for 20 minutes, after which 1.9 Ml of 10 % NaOH was added to stop the reaction. The OD_540_ values were recorded and a standard curve drawn.

The data for the pyruvate standard curve was plotted with pyruvate concentrations on the Y axis and OD_540_ values on the x axis. The slope intercept equation obtained from this graph (y = 21.117x +0.1599) was then used to calculate an extended version of the sodium pyruvate standard curve. This extended version of the pyruvate standard curve was necessary to extrapolate OD_540_ readings for pyruvate concentrations greater than 5 µg/ml. Regression analysis of the linear graph revealed an R^2^ value of.9824. The slope intercept equation obtained from the graph above (y = 21.117x + 0.1599) was used to calculate an extended version of the Sodium Pyruvate Standard Curve. This extended standard curve was utilized in subsequent experiments to convert OD_540_ values to µg of pyruvate. The slope intercept equation for this graph is the following: y = 21.222x + 0.116.

## RESULTS

### *P. aeruginosa* Genome Sequence Analysis for the Presence of L-SDs

*P. aeruginosa* genome sequence analysis reveals the presence of two open reading frames (ORFs) encoding L-SD homologs, *PA2443* and *PA5379* (Figure 9). They are located at position 2742353 to 2740977 and 6056877 to 6058253, respectively (Stover *et al.*, 2000). These ORFs are also referred to as *sdaA* and *sdaB,* respectively.

**Figure 9.**
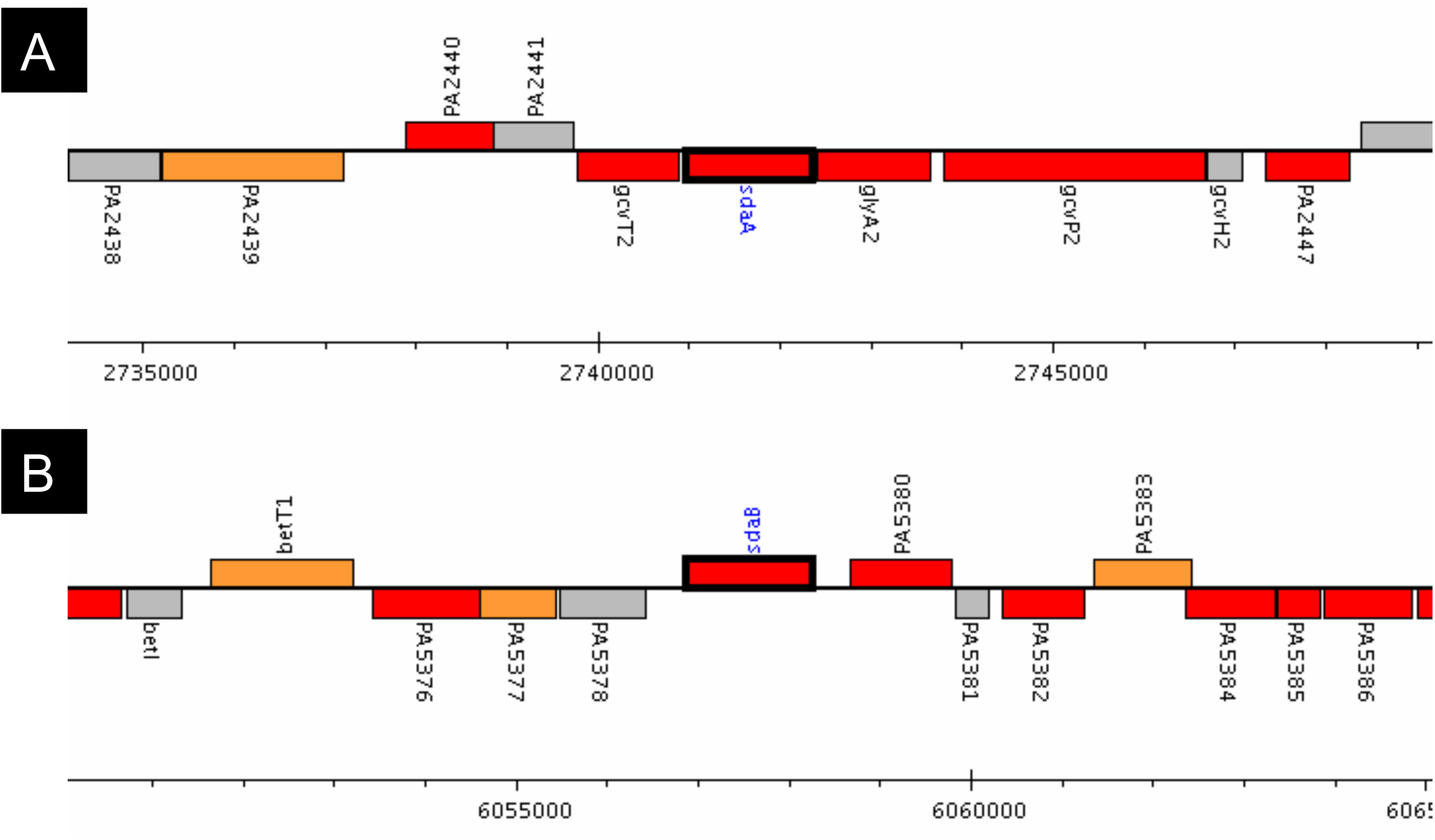
Chromosomal location of *P. aeruginosa sda* genes. s*daA* and *sdaB*. **(A)**. The *sadA* (*PA2443*) gene is found at position 2742353 to 2740977 within the *P. aeruginosa* Strain PAO1 chromosome (http://v2.pseudomonas.com/getAnnotation.do?locusID=PA2443). **(B).** *P. aeruginosa* SdaB is encoded by *sdaB* (*PA5379*) found at position 6056877 to 6058253 within the PAO1 chromosome (http://v2.pseudomonas.com/getAnnotation.do?locusID=PA5379).

The sequence analysis around *sdaA,* revealed the presence of a possible promoter start site between *PA2449-PA2447* with a score of 0.92 on a scale in which the maximum is one and a cut off of 0.8 is deemed an accurate prediction of promoter sequence credibility (Reese, 2001). According to this bioinformatic analysis, the minimal promoter region encompasses the following sequence 5’ CTTGGGA<u>TTGACC</u>AGTTGCAGGTGGGTAA<u>TCGAAT</u>GGGCG<u>A</u>TGCCGCTGT …3’ with the −35 and −10 recognition sequences being 5’ TTGACC 3’ and 5’ TCGAAT 3’, respectively. This promoter region is found upstream of four genes. The analysis of the intergenic region between *sdaA* and its downstream neighboring gene *gcvT2*, revealed the presence of an inverted repeat followed by a string of uracil (5’ACC<u>GCGCCG</u>CTGCGC<u>CGGCGC</u>*TTCTTTTTTCTTT* 3’; underline indicates the inverted repeat) denoting Rho-independent transcriptional termination (Roberts, 1969). The *sdaA* gene appears to be the last gene in a large operon in which the promoter region is found four genes upstream and transcriptional termination occurs immediately after *sdaA* (Figure 9A).

The sequence analysis around *sdaB* revealed the presence of a possible promoter region (Score = 0.69), 744-bp upstream of *sdaB* (Reese, 2001). Thus, the predicted *sdaB* promoter region is localized between 6,056,088 bp and 6,056,133 bp (5’ CTCCAGG<u>TTGGCG</u>CGCAGTGTCTCGACCG<u>TACCTT</u>TCTCGG CGTACTGCT 3’) with the −35 and −10 recognition sequences being 5’ TTGGCG 3’ and 5’ TACCTT 3’, respectively. Analysis of the intergenic region between *sdaB* and its downstream ORF *PA5380*, revealed the presence of an inverted repeat followed by a string of uracil at position 6058497 to 6058520 namely, 5’ CCG<u>GGCT</u>AA<u>AGCC</u>TTCCGG C*TTTTTT* 3’. Thus, *sdaB* appears to be a single gene operon (Figure 9B).

### Analysis of SdaA and SdaB

The *sdaA* and *sdaB* genes encode proteins of 48.9 kD and 49.2 kD, respectively (Table 2). The hydrophobicity plots of the proteins suggest the presence of very hydrophobic regions (Figures 10 and 11). SdaA and SdaB exhibit four and five regions of 20 or more hydrophobic amino acids between amino acids 275 – 375 and 275 - 425, respectively. The hydrophobic nature of these amino acid segments may render them membrane spanning domains, or more likely constituents of the stabilizing hydrophibic core of cytoplasmic L-SD.

**Table 2.** SdaA and SdaB protein analysis.

| **Analysis<sup>a</sup>** | **SdaA\*** | **SdaB\*** |
| --- | --- | --- |
| **Molecular Weight** | 48,945.34 | 49,217.49 |
| **Length** | 458 | 458 |
| **1 microgram = (picomoles)** | 20.431 | 20.318 |
| **Molar Extinction coefficient** | 29,790 ± 5 % | 42,210 ± 5 % |
| **1 A (280) = (mg/ml)** | 1.64 | 1.17 |
| **Isoelectric Point** | 6.61 | 5.75 |
| **Charge at pH = 7** | -1.83 | -8.44 |

\*The SdaA (PA2443) and SdaB (PA5379) protein sequences were obtained from Pseudomonas.com and analyzed using Protean LaserGene DNASTar 2004.

**Figure 10.**
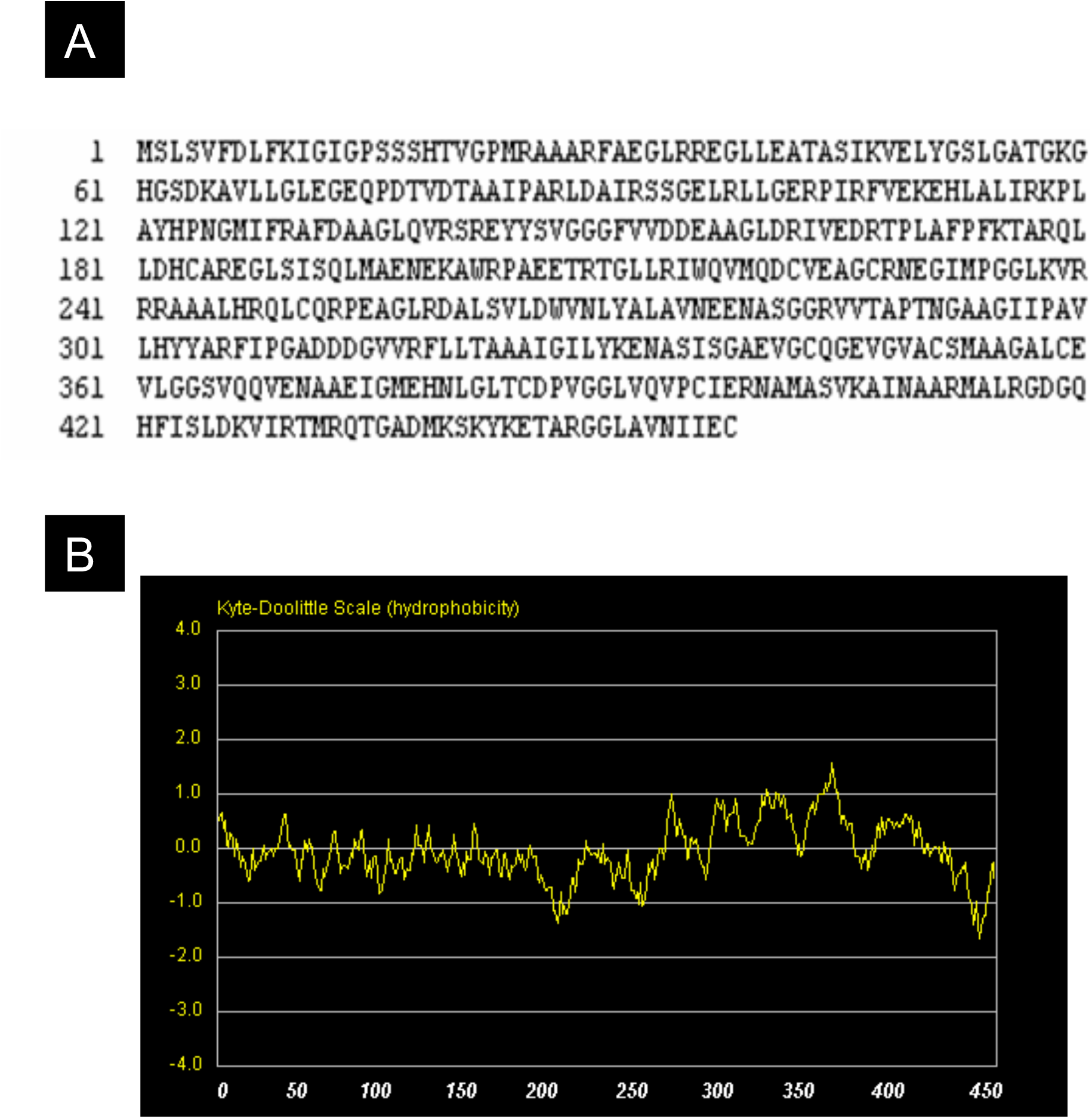
SdaA protein analysis. The amino acid sequence of SdaA **(A)** was analyzed via a Kyte Doolittle Hydrophobicity plot **(B)** ran by Colorado State University http://www.vivo.colostate.edu/molkit/hydropathy/index.htm.

**Figure 11.**
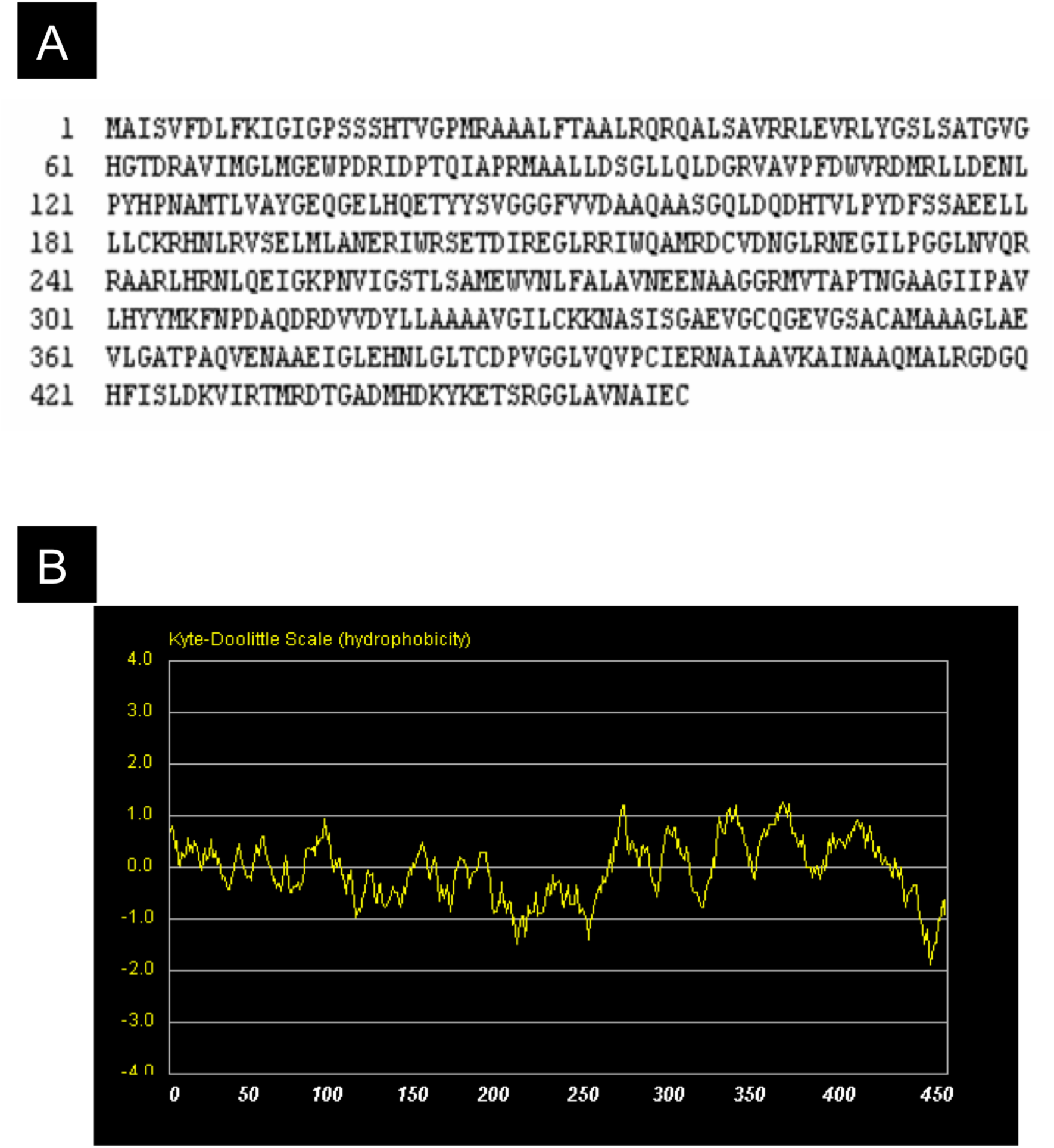
SdaB protein analysis. The amino acid sequence of SdaB (A) was analyzed via a Kyte Doolittle Hydrophobicity plot (B) ran by Colorado State University http://www.vivo.colostate.edu/molkit/hydropathy/index.htm.

The *P. aeruginosa* L-SDs were compared with each other and other L-SD homologs (Table 3). The two *P. aeruginosa* L-SDs, SdaA (EC 4.2.1.13) and SdaB (EC 4.2.1.13), share 62 % homology with one another. *P. aeruginosa* SdaA and SdaB share greater than 41.9 % and 44.4 % homology, respectively, with all other L-SDs analyzed in this study. The least homology was seen with *S. pyogenes* SdaA (12 % with SdaA and 11. 1 % with SdaB). *P. aeruginosa* SdaA also shares 51.3 % and 51.6 % homology with *E. coli* SdaA and SdaB, respectively (Table 3). *P. aeruginosa* SdaB shares 51.8 % and 52.5 % homology with *E. coli* SdaA and SdaB, respectively (Table 3). Both SdaA and SdaB show high homology to L-SDs from *Mycobacteria*. When clustal method analysis was used to analyze the relative similarity of the L-SD enzymes the two *P. aeruginosa* and the two *Mycobacterium* enzymes were most closely related to their congeners (Figure 12). The *E. coli* and *Y. pestis* enzymes were most closely related to each other. *S. pyogenes*, the only Gram positive bacterium in the study was used on the outgroup (Figure 12).

**Table 3.** Comparison of *P. aeruginosa* SdaA and SdaB proteins to its homologs

|  |  |  | Percent Similarity |  |  |  |  |  |  |  |  |
| --- | --- | --- | --- | --- | --- | --- | --- | --- | --- | --- | --- |
|  |  |  | 1 | 2 | 3 | 4 | 5 | 6 | 7 | 8 | 9 |
|  |  |  | *E. coli* | *P. aeruginosa* | *M. tuberculosis* | *C. jejuni* | *E. coli* | *P. aeruginosa* | *S. pyogenes* | *Y. pestis* | *M. bovis* |
|  |  |  | SdaA | SdaA | SdaA | SdaA | SdaB | SdaB | SdaA | SdaA | SdaA |
| D<br>i<br>v<br>e<br>r<br>g<br>e<br>n<br>c<br>e | 1 | *E. coli* SdaA | *** | 51.3 | 48.0 | 42.3 | 76.9 | 51.8 | 9.6 | 84.1 | 48 |
|  | 2 | *P. aeruginosa* SdaA |  | *** | 54.4 | 41.9 | 51.6 | 62.0 | 12.0 | 51.5 | 54.4 |
|  | 3 | *M. tuberculosis* SdaA |  |  | *** | 39.6 | 47.7 | 52.6 | 12.0 | 47.7 | 100 |
|  | 4 | *C. jejuni* SdaA |  |  |  | *** | 42.5 | 40.7 | 12.3 | 42.5 | 39.6 |
|  | 5 | *E. coli* SdaB |  |  |  |  | *** | 52.5 | 12.0 | 76 | 47.7 |
|  | 6 | *P. aeruginosa* SdaB |  |  |  |  |  | *** | 11.1 | 51.5 | 52.6 |
|  | 7 | *S. pyogenes* SdaA |  |  |  |  |  |  | *** | 10.2 | 11.7 |
|  | 8 | *Y. pestis* SdaA |  |  |  |  |  |  |  | *** | 47.1 |
|  | 9 | *M. bovis* SdaA |  |  |  |  |  |  |  |  | *** |

Clustal analysis was done using *P. aeruginosa* SdaA and SdaB, *E. coli* and SdaB, *M. tuberculosis* SdaA, *M. bovis* SdaA, *C. jejuni* SdaA, *Y. pestis* SdaA and *S. pyogenes* SdaA.

**Figure 12.**
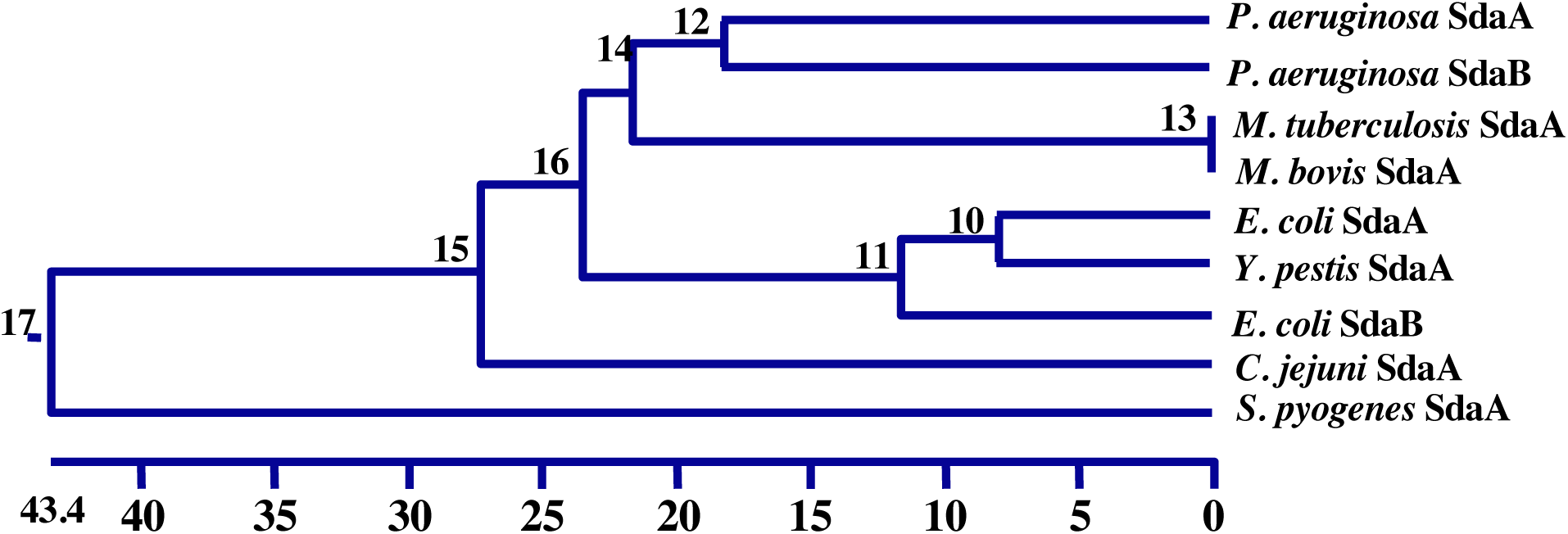
Clustal analysis of LSD homologs. This analysis revealed that the two *P. aeruginosa* L-SDs and the two *Mycobacterium* enzymes were most closely related. *E. coli* and *Y. pestis* enzymes were most closely related. *C. jejuni* is sister to these species and *S. pyogenes*, being the only Gram positive bacterium in the study is separated from all other species in the study

### Minimal Plating Experiments

Strains PAO1, PAO*sdaA*, and PAO*sdaB* were plated on NIV minimal media with various carbon sources: glucose (NIV-Glu), serine (NIV-Ser), glycine (NIV-Gly), leucine (NIV-Leu), serine-glycine (NIV-Ser-Gly), serine-leucine (NIV-Ser-Leu), serine-glycine-leucine (NIV-Ser-Gly-Leu). As controls the cultures were plated on NIV and NMM media. Both the number of the colonies and the characteristics were observed. The PAO1 colonies grew best in NIV-Glu (the positive control) media with the colony size recorded at 4.0 mm (Table 4). These colonies were large, healthy, pronounced, and harbored a large volume of cells per colony. As expected no colonies were observed in NMM media that served as our negative control. We expected no growth in the NIV plate that is also considered minimal media, but contains the amino acids isoleucine and valine. However, there were colonies, which were small (1 mm) and translucent with 91 % survival (Table 4). Similar translucent colonies were also observed on NIV-Ser (Table 4). The colony size on NIV-Ser-Leu was also 1 mm but the colonies were tight, compact, much more pronounced, and the amount of cells/ colony was visibly greater (10x) than that of the PAO1 cells grown on NIV media alone and at least five-fold more than NIV-Ser media. A significant decrease in colony % survival (78 %) was noted for PAO1 colonies in NIV-Ser media (Table 4). An even greater significant change in colony survival (63 %) was observed in NIV-Ser-Leu media (Table 4).

**Table 4.** Colony size and percentage survival of PAO1 and its isogenic *sda* mutants on various media.*

| **Media<sup>a</sup>** | **PAO1** |  | **PAO*sdaA*** |  | **PAO*sdaB*** |  |
| --- | --- | --- | --- | --- | --- | --- |
|  | **Size (cm)** | **%** | **Size (cm)** | **%** | **Size (cm)** | **%** |
| **Glucose** | 4 | 100 | 4.5 | 100 | 4 | 100 |
| **Ser-Gly-Leu** | 3.5 | 89 | 2 | 85 | 3 | 61 |
| **Ser-Gly** | 3.5 | 88 | 1.5 | 73 | 3 | 81 |
| **Ser-Leu** | 1 | 63 | 0.8 | 78 | 1 | 48 |
| **Serine** | 1 | 78 | 0.5 | 79 | 1 | 71 |
| **Glycine** | 2.5 | 87 | 1.3 | 80 | 2.5 | 64 |
| **Leucine** | 3 | 63 | 1 | 74 | 4 | 50 |
| **NIV** | 1 | 91 | 0.5 | 87 | 1 | 62 |
| **NMM** | 0 | 0 | 0 | 0 | 0 | 0 |

\*Strains PAO1, PAO*sdaA*, and PAO*sdaB* were plated on differential media. Colony size and percentage colony survival was recorded after 3 days of growth. NIV – Newman Isoleucine-Valine Media; NMM – Newman minimal media

The number of the colonies in NIV-Ser-Gly was similar to NIV-Glu but the colony characteristics were distinctly different. The colonies on NIV-Gly media (2.5 mm) were translucent with visibly fewer cells per colony than in NIV-Ser-Gly (Table 4). The estimated amount of cells/ colony in NIV-Gly is 1/10^th^ the NIV-Glu.

The best growth was observed in NIV-Ser-Gly-Leu with colonies slightly smaller (3.5 mm) compared to those in NIV-Glu media (Table 4). The percentage of PAO1 colonies that formed on NIV-Ser-Gly-Leu and NIV-Ser-Gly plate media as compared to those that formed on NIV-Glu media is ∼ 89 suggesting failure of 10 % of colonies to grow on this media (Table 4).

### PAO*sdaA* Colony Description

The mutant strain PAO*sdaA* shared almost similar percentage survival as the wild type PAO1. However, the colony size is smaller in all the media tested (Table 4). PAO*sdaA* exhibited similar growth properties as compared with PAO1 in NIV-Glu media. The *sdaA* mutant also does very poorly on NIV-Ser (Table 4).

The percentage survival in NIV-Ser-Gly-Leu was comparable to that for PAO1 but the colonies were tight, compact, and healthy (2.5 mm). The only observable difference was the decrease in colony size, attributable of course to the absence of one of the L-SD encoding genes (*sdaA*). A significant decrease in colony size was also observed when PAO*sdaA* cells grown in NIV-Ser-Gly (1.5 mm), NIV-Ser-Leu (0.75 mm) and NIV-Ser (0.5 mm) as compared to PAO1 (Table 4).

The presence of trace amounts of leucine appears to enhance the growth of strain PAO*sdaA.* The survival for strains growing on NIV-Ser-Leu (78 %) and NIV-Leu (74 %) were significantly greater than PAO1 cells grown on NIV-Ser-Leu (63.33 %) and NIV-Leu (73.78 %) (Table 4).

### PAO*sdaB* Colony Description

The mutant strain PAO*sdaB* showed similar colony morphology as that of parent strain PAO1 (Table 4). There is an overall decrease in percentage survival in all the media tested.

A noticeable reduction was observed in NIV-Ser-Gly-Leu (61.48 %) (Table 4). The colonies on this media were also tight, compact, and healthy. Of significant note is the fact that unlike PAO*sdaA* colonies that appeared to benefit slightly by leucine supplementation of NIV-Ser media, no increase in colony size was observed between PAO*sdaB* grown on NIV-Ser-Gly-Leu (3.5 mm) versus those grown on NIV-Ser-Gly (3.5 mm) (Table 4).

Similar colony size was observed when PAO*sdaB* colonies were grown on NIV-Ser and NIV-Ser-Leu. However, a significant decreased percentage survival was observed in NIV-Ser-Leu media versus NIV-Ser media (48 % vs 71 %) and also in NIV-Ser-Gly-Leu versus NIV-Ser-Gly media (61 % vs 81 %) (Table 4). The presence of trace amounts of leucine appears to inhibit growth of strain PAO*sdaB.*

### Growth Curve Properties of PAO1

*P. aeruginosa* PAO1 growth in rich liquid media and in minimal media supplemented with various amino acids was analyzed. From Figure 13 the growth curves, the doubling time was calculated. As expected PAO1 growth was optimal in rich LB media (Figure 13) with a doubling time of approximately 37.4 minutes (Figure 13, Table 5). PAO1 was also able to grow in NIV-Glu minimal media with a doubling time of 80.5 min. In the presence of Ser-Gly and Ser-Gly-Leu the growth rate was drastically reduced with a doubling time change of 285 and 210 min, respectively (Figure 13, Table 5). PAO1 was unable to utilize serine and leucine as the sole carbon source (Figure 13).

**Table 5.** Doubling Times of PAO1 and L-SD Mutants.

| Strains<sup>a</sup> | Doubling Times<sup>b</sup> (min) |  |  |  |
| --- | --- | --- | --- | --- |
|  | LB | NIV | NIV | NIV |
|  |  | Glu | Ser-Gly-Leu | Ser-Gly |
| **PAO1** | 37.4 | 80.5 | 210.2 | 285 |
| **PAOsdA** | 38.5 | 82 | 245.3 | 811.9 |
| **PAOsdB** | 37.8 | 80.9 | 265 | 681.8 |

a. The L-serine deaminase mutants, PAOsdA and PAOsdB, are isogenic with PAO1 containing (ISphoA/hah) insertions at the sdaA and sdaB loci respectively.
b. *P. aeruginosa* was grown overnight in glucose minimal media and subcultured in minimal media with serine and amino acid supplements. The OD600 was recorded every hour for 16 hours and the doubling time calculated.

**Figure 13.**
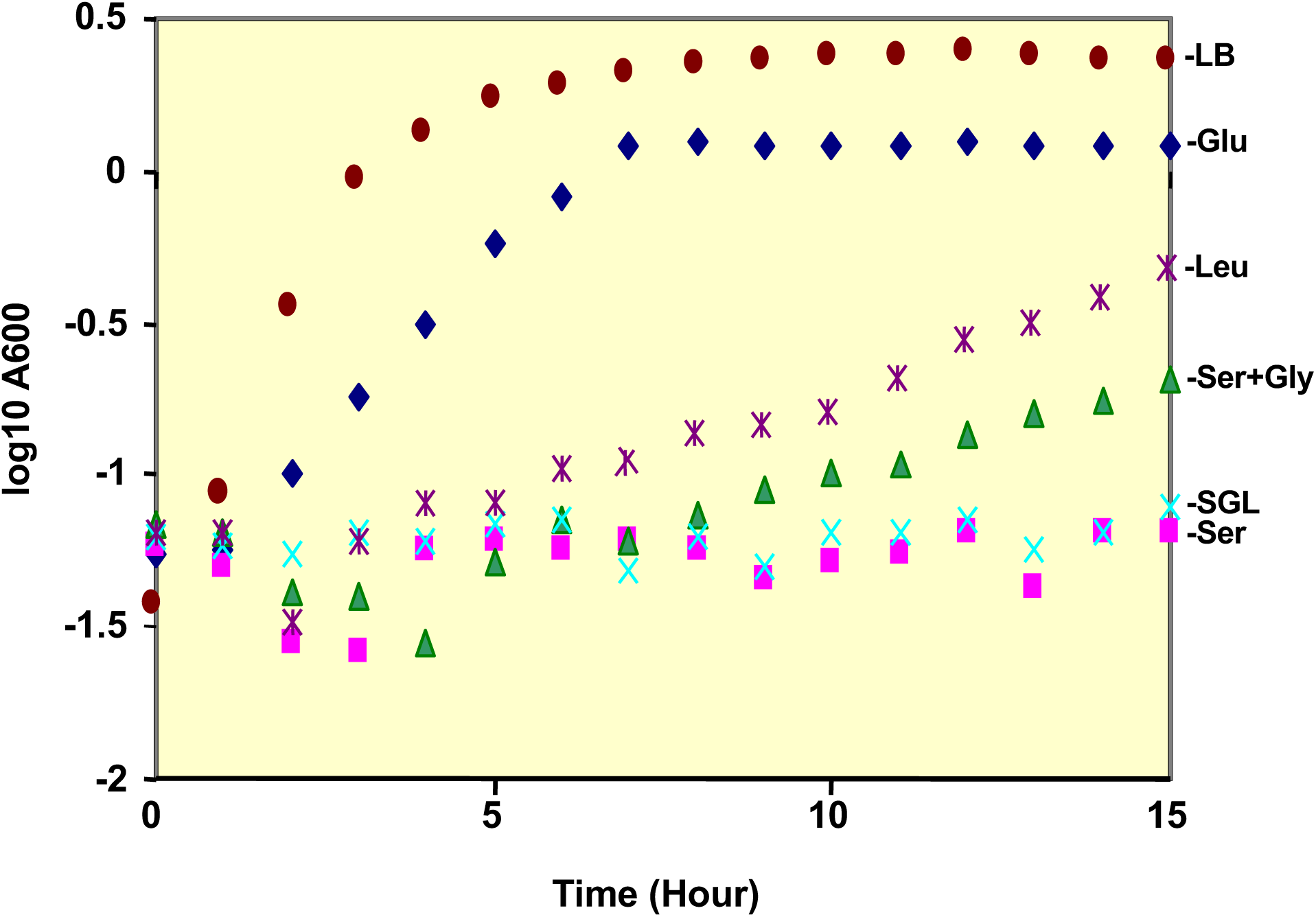
*P. aeruginosa* PAO1 growth in various media. PAO1 growth was optimal in rich LB media 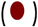, NIV media with Glucose (♦), serine (▮), serine and glycine (▴), serine and leucine (X), serine, glycine and leucine 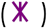. The growth OD600 was determined for every hour for 15 hours.

### Growth Properties of *sdaA* and *sdaB* Mutants

The growth properties of the L-SD mutants, PAO*sdaA* and PAO*sdaB* were compared with the parental PAO1. The growth properties of PAO*sdaA* and PAO*sdaB* did not differ significantly from PAO1 in LB media (Figure 14A, Table 5) and NIV-Glu (Figure 14B, Table 5). Similar to PAO1, the mutants were able to grow on Ser-Gly-Leu, however the doubling time was slightly reduced suggesting a slight increase in growth rate (Figure 14F, Table 5). The growth of both *sdaA* and *sdaB* mutants was significantly compromised in NIV-Ser-Gly media that lacked leucine. Where PAO1 growth doubled at 285 min, the *sdaA* and *sdaB* mutants doubling times were significantly longer, 812 and 682 min, respectively (Figure 14E, Table 5).

**Figure 14.**
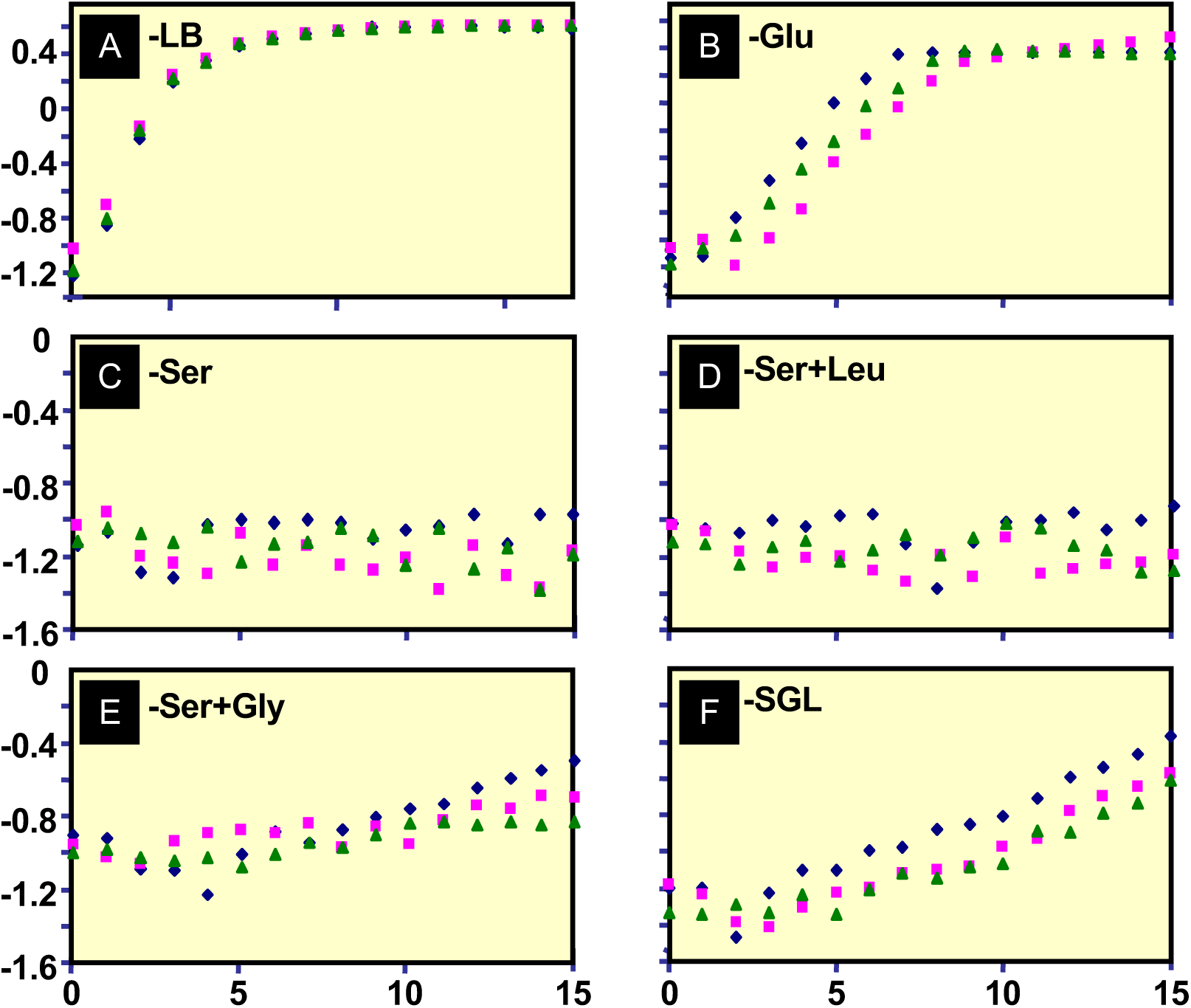
Growth characteristics of *P. aeruginosa sda* mutants. The growth pattern of PAO*sdaA* (◼) and PAO*sdaB* (▴) were compared with their isogenic parent strain PAO1 (♦). The cells were cultured in **(A)** LB, **(B)** glucose, **(C)** NIV minimal media with serine, **(D)** NIV minimal media with serine and leucine, **(E)** NIV minimal media serine and glycine, and **(F)** NIV minimal media with serine, glycine and leucine.

As expected, similar to the prototype strain PAO1, both PAO*sdaA* and PAO*sdaB* mutants failed to grow in NIV with serine as the sole carbon source (Figure 14C, Table 5). In fact, the addition of leucine (Figure 14D, Table 5) to NIV-Ser failed to support growth. However, addition of glycine to NIV-Ser media did promote growth of both mutants (Figure 14E, Table 5).

### *P. aeruginosa* L-SD Activity

The L-SD enzyme assay was performed on strain PAO1 grown in NIV-Glu-Gly and NIV-Ser-Gly-Leu. In both media, there is a lag of 20-30 min, followed by a quick burst of activity that plateaus at 80 min (Figure 15). The L-SD activity rate in Ser-Gly-Leu (0.155 µg/ min) is twofold higher than the activity rate in NIV-Glu-Gly media (0.076 µg/ min).

**Figure 15.**
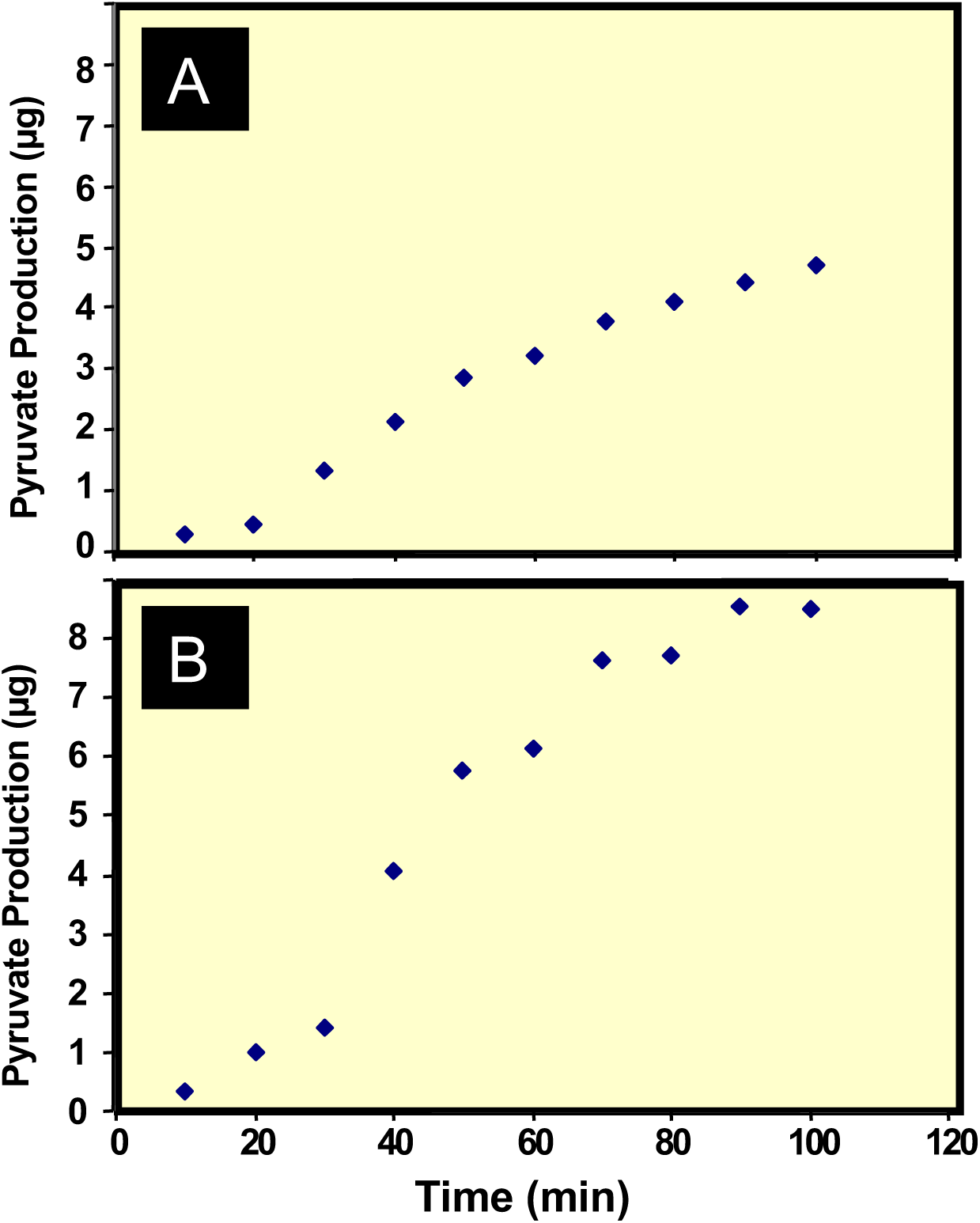
The L-serine deaminase enzyme activity of *P. aeruginosa* PAO1. The L-SD activity was measured after culturing the bacteria in NIV media with **(A)** glucose and glycine (NIV Glu-Gly) and, **(B)** serine, glycine and leucine (NIV Ser-Gly-Leu).

After determining the time of maximal activity, subsequent assays were done on 14 hr cultures, and the amount of pyruvate made was determined after 50 min. The L-SD activity of PAO1 in NIV-Ser-Gly-Leu, NIV-Glu-Gly, NIV-Glu, and NIV-Gly-Leu were analyzed (Table 6). There is a significant 28.6 % decrease in L-SD activity upon addition of leucine to the NIV-Glu media (Table 6). In contrast, a 3.2-fold increase in L-SD activity was observed in NIV-Ser-Gly-Leu when compared to NIV-Glu-Gly media (Table 6). No significant difference in L-SD activity was observed in PAO1 grown in NIV-Gly-Glu media or NIV-Glu media (Table 6).

**Table 6.** L-serine deaminase activity of *sda* mutants.

| Strains<sup>a</sup> | L-Serine Deaminase Activity<sup>b</sup> |  |  |  |
| --- | --- | --- | --- | --- |
|  | Ser-Gly-Leu | Glu | Glu-Gly | Glu-Leu |
| **PAO1** | 6.8 | 2.1 | 2.4 | 1.5 |
| **PAO*sdaA*** | 5.2 | 1.8 | 1.9 | 1 |
| **PAO*sdaB*** | 2.7 | 1.7 | 2.4 | 0.8 |

a. The L-serine deaminase mutants, PAO*sdaA* and PAO*sdaB*, are isogenic with PAO1 containing insertions of (ISphoA/hah) in each loci.
b. L-Serine-Deaminase activity was measured after 14 hr growth on NIV media with serine, glycine and leucine (Ser-Gly-Leu), with glucose (Glu), glucose and glycine (Glu-Gly) and glucose and lysine (Glu-Leu). The activity is expressed as μg of pyruvate produced

When strain PAO*sdaA* was grown in the same media the same general pattern of induction was observed, however the amount of pyruvate produced is generally lower than the parent PAO1 (Table 6). No observable increase in activity was observed when glycine was added to NIV-Glu media (Table 6). A 44% decrease in activity was observed when leucine was added to NIV-Glu media and a 2.9-fold increase in L-SD activity was observed between strains grown in NIV-Ser-Gly-Leu as compared to NIV-Glu media (Table 6).

In PAO*sdaB* a decreased pyruvate production was observed as compared to PAO1 in NIV-Glu, NIV-Ser-Gly-Leu, and NIV-Gly-Leu media (Table 6). However, in NIV-Glu-Gly media PAO*sdaB* and PAO1 showed indistinguishable L-SD activity. A 1.4-fold increase in activity was observed when glycine was added to NIV-Glu media (Table 6). A 1.6-fold increase in L-SD activity was observed in Ser-Gly-Leu media as compared to NIV-Glu media and a 53 % decrease in L-SD activity was observed upon leucine supplementation (Table 6).

### Serine Growth Mutants Characteristics

After growth for 36 hours the OD _600_ values for the five serine utilizing mutants (*ser1* to *ser5)* were recorded and their L-SD activity was assayed (Table 7). Unlike the parent PAO1, *ser1, ser2, ser 3, ser4* and *ser5* were able to utilize serine as a sole carbon source (Table 7).

**Table 7.** L-serine deaminase activity of serine growth (ser) mutants.

| Strains<sup>a</sup> | OD<sub>600</sub><sup>b</sup> | L-Serine Deaminase Activity<sup>c</sup> |  |
| --- | --- | --- | --- |
|  |  | Ser-Gly-Leu | NIV |
| **PAO1** | 0.014 | 6.8 | N/A |
| **PAOsdA** | 0.058 | 5.2 | N/A |
| **PAOsdB** | 0.02 | 2.7 | N/A |
| **Ser1** | 0.501 | 5.9 | 1.2 |
| **Ser2** | 0.323 | 6.1 | 1.7 |
| **Ser3** | 0.176 | 8 | N/A |
| **Ser4** | 0.744 | 6.3 | 1.7 |
| **Ser5** | 0.36 | 7.9 | 1.6 |

a. The serine growth mutants, *ser1* to *ser 5*, that can utilize serine, were isolated by continuous restreaking of strain PAO1 on minimal media plates with serine as the sole carbon source.
b. Strains were grown for 36 hr on NIV media with serine as the sole carbon source at which point the OD<sub>600</sub> value was recorded.
c. L-Serine deaminase activity was measured after 14 hr growth on NIV alone and NIV media with serine, glycine and leucine (Ser-Gly-Leu). The activity is expressed as μg of pyruvate produced after 50 minute incubation as described in Materials and Methods.

## DISCUSSION

All pathogens require high energetic influxes to counterattack the host immune system and without this energy bacterial infections are easily cleared. This study was an investigation into a highly bioenergetic pathway in *Pseudomonas aeruginosa* involving the amino acid L-serine and the enzyme L-serine deaminase (L-SD). Recent evidence linked L-SD directly to the pathogenicity of several organisms including but not limited to *Campylobacter jejuni, Mycobacterium bovis*, *Streptococcus pyogenes*, and *Yersinia pestis* (Velayudhan *et al.*, 2004). We hypothesized that *P. aeruginosa* L-SD is likely to be critical for its virulence. We analyzed the ability of *P. aeruginosa* to utilize serine and the role of SdaA and SdaB in serine deamination by comparing mutant strains of *sdaA* (PAO*sdaA*) and *sdaB* (PAO*sdaB*) with their isogenic parent *P. aeruginosa* PAO1.

### *P. aeruginosa* Harbors Two L-SDs

*P. aeruginosa* Genome sequencing revealed the presence of two genes encoding L-SDs, *sdaA* and *sdaB* (*PA2443* and *PA5379*) (Stover *et al.*, 2000). These two *P. aeruginosa* L-SDs, SdaA and SdaB share 62 % homology to one another (Table 3). The high homology suggests that one of these two genes became incorporated into the genome via duplication of the existing *sdaA* or *sdaB*. The dissimilarity between these two proteins is also large enough to suggest that the duplication event took place early in the evolution of *P. aeruginosa*.

Multi-copy genes are usually essential genes, encoding a product that the bacterium cannot afford to lose. They offer protection against mutation and loss of phenotype. L-SD is usually encoded multiple times within a bacterial genome. *P. aeruginosa* harbors two genes encoding L-SD (Stover *et al.*, 2000). *E. coli* bears three copies of L-SD in its genome (Keseler *et al.*, 2005). The presence of multiple copies of this enzyme in bacterial genomes adds weight to the hypothesis that L-SD is important for growth of a bacterium.

In *E. coli* and *P. aeruginosa,* each multicopy L-SD was proven to be functional and yet both bacteria exhibited growth failure with serine as the sole carbon source (Alfoldi & Rasko, 1970). Though the enzyme is encoded multiple times in the genome and its expression should be high within the constitutive cell, no growth is observed in *E. coli* unless leucine is added to the media, and no growth is observed in *P. aeruginosa* unless glycine is added to the media (Newman *et al.*, 1985)(Table 4, 5). In *E. coli* it is known that L-SD is kept under tight negative regulation by the Lrp protein and when leucine is added this negative inhibition is released and growth is allowed (Newman *et al.*, 1985). Although an Lrp homolog is present in the *P. aeruginosa* genome, the opposite phenotypic effect is observed when leucine is added to the media (Table 4, 5, 6).

The role of glycine in L-SD induction is novel in the bacterial literature. In subsequent sections, the possibility of glycine enhancing transport of L-serine into the cell is discussed as a possible explanation of the induction potential of glycine on L-SD expression.

### *P. aeruginosa* L-SDs are Nonessential

The loss of *sdaA* and *sdaB* had no effect on *P. aeruginosa* growth in LB media (Figure 14A). This is not surprising as LB contains carbon sources in the media other than serine that allow growth and therefore *sdaA* and *sdaB* were not essential. In addition, the absence of the L-SD encoding genes did not affect growth in glucose minimal media (Figure 14B). This also makes sense given the fact that the carbon source utilized for growth in this media is glucose and not L-serine. In fact, as expected, the L-SD activity in LB is lower than in minimal medium with glucose (Table 6). It is possible that the use of serine becomes critical at stationary phase when the nutrients in the media have been exhausted. The assay was performed after 14 hours of growth where in rich LB media the culture has passed the stationary phase and has entered the death phase (Figure 14A). It is possible that at this stage of growth, PAO1 L-SD activity will be reduced, especially if as in other organisms L-serine is the first amino acid utilized (Prüß, 1994). Thus, a detailed time course experiment is required to determine the L-SD activity.

### *P. aeruginosa* cannot utilize Serine as a Sole Carbon Source

PAO1 colony size on the minimal plating experiments (Table 4), suggests that *P. aeruginosa* is not capable of growth with serine as a sole carbon source This finding is further confirmed by growth curves (Figure 14C). Like the prototype strain, PAO1, no growth was observed in both the *sdaA* and *sdaB* mutant strains in media with L-serine as the sole carbon source (Figure 14C). *E. coli* also exhibits no growth on serine as the sole carbon source, but as stated earlier the enzymes, though functional, are held under tight negative Lrp regulation.

In this study, we have also isolated five serine growth mutants (Table 7) that now can utilize serine as the sole carbon source. Likely these strains have mutations that either up regulate an activator or have inactivated a repressor. In either case, the net result is constitutive expression of L-SD allowing the bacteria to utilize serine. Future mapping of the mutations in these strains will likely reveal the intricate mechanisms involved in regulating the expression of L-SDs.

### Both SdaA and SdaB Contribute to the Total L-SD Activity

The similarity between both *P. aeruginosa* L-SDs and the L-SDs of *C. jejuni, M. bovis, Y. pestis*, and *E. coli* suggests that these enzymes work through a similar mechanism in these bacteria and likely have a similar function (Table 3). Our analysis of PAO*sdaA* and PAO*sdaB* mutants show that both SdaA and SdaB contribute to the total L-SD activity (Table 6). The loss of either *sdaA* or *sdaB* in serine-glycine media contributed to a strikingly large increase in doubling time as compared to PAO1, suggesting again that both SdaA and SdaB are contributing to PAO1 growth in glycine supplemented serine minimal media (Table 5). Based on the ability of the mutants to convert serine to pyruvate, we estimated that SdaA and SdaB contribute 34 % and 66 % of the total activity, respectively (Table 6). Hence, loss of one gene is unable to abolish the activity completely. Since SdaB appears to be the major contributing L-SD, we predict that PAO*sdaB* is likely to be less virulent as compared to PAO*sdaA.*

### Glycine Enhances *P. aeruginosa* Serine Utilization

*P. aeruginosa* cannot grow on serine minimal media alone (Figure 14C). However, the addition of glycine resulted in a colony size similar to the one obtained with glucose media. *P. aeruginosa* is also capable of growth in minimal media with glycine as the sole carbon source (Table 4). However, the extent of glycine utilization was minimal in comparison to PAO1 growth in serine media supplemented with glycine (Table 4). The glycine-dependent growth in serine media can also been seen in the growth curve (Figure 14E). These data suggest an ability of L-glycine to enhance L-serine utilization in *P. aeruginosa* PAO1. This is supported by the total L-SD activity where the activity was elevated in media with serine and glycine as compared to growth in glucose minimal media alone (Table 6).

The analyses of the *sdaA* and *sdaB* mutant strains suggest that the SdaA L-SD activity is glycine-dependent (Table 6). The prototypic PAO1 that harbors active SdaA and SdaB showed a 2.8-fold induction in L-SD activity upon addition of glycine to serine media (Table 6). Strain PAO*sdaB* that has an active SdaA exhibited a similar induction (2.7- fold) in the presence of L-serine (Table 6). In contrast, PAO*sdaB* that harbors active SdaA showed no significant induction of L-SD activity upon addition of L-serine (Table 6). Interestingly, glycine induction of L-SD activity in glucose media was observed in strain PAO*sdaB* (Table 6). No effect was observed in PAO1 or in PAO*sdaA* under the same experimental conditions (Table 6).

It is not known how L-serine enters the *P. aeruginosa* cell. Genomic analysis reveals the absence of specific L-serine membrane transporters (Stover *et al.*, 2000). We speculate that an L-serine / L-glycine symporter is present within the *P. aeruginosa* outer membrane which allows L-serine entrance into the cell only in the presence of L-glycine. If this were the case then in the presence of glycine, serine would enter the cell and induce transcription of the L-SD enzymes. Indeed 4-fold induction of L-SD activity was observed when glycine and serine were present in the media (Table 6). If glycine were not present, then serine would not be able to enter the cell, L-SD expression would not be induced, and L-SD activity would remain constitutive. Indeed, no significant difference in L-SD activity was observed upon addition of glycine to glucose minimal media (Table 6).

### Serine is the Inducer of *P. aeruginosa* L-SD Activity

Our data also shows that glycine alone cannot induce the L-SD activity as there was no significant increase in L-SD activity between PAO1 grown in glucose and glucose-glycine media (Table 6). Thus, serine has to be present for the induction of L-SD activity. A 3.2-fold increase in total pyruvate production was observed between strain PAO1 grown in serine-glycine as compared to glucose media (Table 6). This strongly argues that L-serine itself is an inducer of L-SD activity. Interestingly, there is a slight increase of SdaB activity in glucose-glycine media in the absence of serine. This might just be a basal activation of *sdaB* expression but full induction requires the inducer serine. The fact that L-serine is an inducer of L-SD makes biological sense. It is bio energetically inefficient to transcribe a gene, and translate a protein, if that protein is not going to be used within the cell.

### Leucine Inhibits *P. aeruginosa* L-SD Activity

In. *E. coli* the addition of L-leucine to serine minimal media induces the production of L-SD and the concomitant breakdown of L-serine for energy requirements (Newman *et al.*, 1985). This induction takes place via regulation of the **l**eucine-**r**esponsive regulatory **p**rotein **Lrp** which negatively regulates *E. coli sda* genes (Calvo & Matthews, 1994; Newman & Lin, 1995). *P. aeruginosa* genome analysis revealed the presence of an *E. coli* Lrp homolog (PA5308)}; Stover *et al.*, 2000). We hypothesized that L-SD regulation in *P. aeruginosa* occurred via the same mechanism.

Supplementing serine with L-leucine both in the minimal plates (Table 4) and liquid media (Figure 14D) failed to support *P. aeruginosa* growth. In contrast, a slight but significant decrease in doubling time (that is increased growth rate) was observed in NIV media supplemented with serine-glycine-leucine as compared to serine-glycine alone (Table 5). This increased growth rate may be due to leucine and not serine catabolism since growth in minimal media with leucine as the sole carbon source was observed (Table 4). In addition, the small differences in the doubling time between PAO1 grown in serine-glycine-leucine with serine-glycine explain why colonies on minimal media plates with similar amino acid combinations appeared indistinguishable (Table 4).

Analysis of the whole cell L-SD assays revealed a 29 % decrease in L-SD activity upon L-leucine addition to glucose media (Table 6). It appears that not only does L-leucine fail to contribute to L-serine deamination in *P. aeruginosa* but it down regulates the total L-SD activity. In fact, both SdaA and SdaB activity were inhibited by the addition of L-leucine to glucose minimal media (Table 6). The relative rates of leucine inhibition showed a slightly greater inhibitory effect on SdaA (53.0 %) than SdaB (44 %) (Table 6). This is in sharp contrast to the L-SD regulatory system in *E. coli,* where leucine interaction with the Lrp protein induces L-SD activity (Newman & Lin, 1995). Thus, it appears that leucine-dependent regulation of *P. aeruginosa* L-SD activity is Lrp-independent.

### Infectivity Study

The hypothesis of this study is that *in vivo* utilization of the amino acid L-serine confers a highly efficient bioenergetic pathway to *P. aeruginosa*, which is utilized to counterattack host immune defense. Since both SdaA and SdaB contribute to the total L-SD activity, a double mutant is required to determine the role of serine deamination in pathogenicity. In addition, the strain *P. aeruginosa* PAO1 is less virulent as compared to *P. aeruginosa* PA14. The second half of this study focused on construction of suicide vectors harboring insertional inactivated gene cassettes of *sdaA* and *sdaB.* The suicide vectors are designed to lack a compatible origin of replication with *P. aeruginosa* (Schweizer & Hoang, 1995). This characteristic forces them to integrate into the *P. aeruginosa* chromosome at regions of homology. Since, the suicide vectors constructed in this study bear the *sdaA*::*aacCI, sdaB*::*aacCI*, and *sdaB*::*aphI* cassettes, it is possible to selectively mutate the *sdaA* and *sdaB* genes in *P. aeruginosa* strains of interest including PA14 (Schweizer & Hoang, 1995).

Subsequent investigations into the role of L-SD in *P. aeruginosa* pathogenicity will require animal model infectivity studies. Aspects of innate immunity are evolutionarily conserved amongst vertebrates, invertebrates, and plants (Kurz & Ewbank, 2003). In response, the evolution of vertebrate, invertebrate, and plant pathogens has followed a similar trend. The result is that bacterial pathogenic mechanisms which are utilized to invade invertebrate hosts, will serve the same or similar functions in vertebrate and plant hosts and genes essential for virulence in invertebrates prove to be essential for virulence in other biological kingdoms (Tan *et al.*, 1999b; Gan *et al.*, 2002; Kothe *et al.*, 2003; Aballay *et al.*, 2000; Garsin *et al.*, 2001; Sifri *et al.*, 2003; Jansen *et al.*, 2002). To study the *in vivo* characteristics of *P. aeruginosa,* several model systems may be employed. Apart from the murine model system, the most commonly used are the *Arabidopsis thaliana*, *Drosophila melanogaster, and Caenorhabditis elegans* infectivity model systems (Rahme *et al.*, 1995; Vodovar *et al.*, 2004; Tan *et al.*, 1999a) The *C. elegans-P. aeruginosa* pathogenicity model system is the simplest model system to study bacterial pathogenicity.

*C. elegans* is a nematode which exhibits a three-day life span and reaches 1.5 mm in length when fully grown (Brenner, 1974; Byerly *et al.*, 1976). It contains rudimentary digestive, nervous, muscle, and reproductive systems (Brenner, 1974). It is entirely transparent and the fate of each of its adult somatic cells has been mapped. *C. elegans* is ubiquitous in the soil, where it feeds on various bacteria in its surroundings. In the lab it is routinely cultured on lawns of the bacterium *E. coli* strain OP50, but if given no alternative, it will eat whichever bacterium is provided for it (Lewis & Fleming, 1995). When fed *P. aeruginosa* Strain PA14, the worms acquire a gut infection, leading to reduced viability of the worms (Tan *et al.*, 1999a). This model system is especially useful for our purposes for two reasons. The first is that the severity of the infection correlates to the virulence of the bacterium (Tan *et al.*, 1999b). The second is that observed variations from the prototype strain by isogenic mutants reveal the contribution of individual genes to the virulence of *P. aeruginosa* (Ausubel, 2005). When fed *P. aeruginosa* strain PAO1, worms are killed via a distinct toxicity mechanism involving the production of cyanide and other virulence factors (Gallagher & Manoil, 2001). This worm-killing assay is useful in determining variations in virulence factor production among *P. aeruginosa* strains. The mutant strains (PAO*sdaA* and PAO*sdaB*) utilized in this study are derived from the prototype strain PAO1. This strain is not sufficiently virulent to establish an infection in the gut of *C. elegans* (Tan *et al.*, 1999a). *P. aeruginosa* PA14 shows increased virulence as compared to strain PAO1 (Tan *et al.*, 1999a) as it is capable of forming a persistent infection in the gut of *C. elegans* (Mahajan-Miklos *et al.*, 1999). The PA14-derived L-SD mutants will facilitate further elucidation of the role of L-SD in *P. aeruginosa* metabolism and pathogenicity.

### *P. aeruginosa* L-SD and its role in pathogenesis

A pathogenic organism such as *P. aeruginosa* needs to obtain energy from host metabolites. Given the selective advantage conferred on organisms with high L-SD activity *in vitro*, the ability to utilize L-serine released by proteolytic cleavage should confer a selective advantage on these pathogenic organisms *in vivo* (Zinser & Kolter, 2000). This study showed the ability of *P. aeruginosa* to break down the amino acid L-serine via two enzymes SdaA and SdaB (Table 7). The induction of L-SD activity requires L-glycine (Table 7). *In vivo* and during infection, *P. aeruginosa* will be surrounded by a large enough concentration of amino acids including glycine which will induce the activity of L-SD. This increased L-SD activity will be utilized by *P. aeruginosa* to obtain energy. The contribution to L-SD activity by SdaA and SdaB will likely depend on the ratio of glycine, serine and leucine in the surrounding tissues, because differential induction intensities were observed between the two L-SD enzymes (Table 7).

In the presence of so many other carbon sources, why should the breakdown of one amino acid prove to be so important? L-SD was proven to be essential for infectivity by *C. jejuni* and *M. bovis* and thus it may also be required for *P. aeruginosa* infectivit*y* (Velayudhan *et al.*, 2004; Chen *et al.*, 2003)*. P. aeruginosa*, is known for its metabolic diversity, with a capacity to utilize over 200 different carbon sources for energy (Vasil, 1986). It has a vast amount of metabolic genes and metabolic regulation assisting in its energy acquisition capabilities (Vasil, 1986). If further investigation, proves L-SD to be important for pathogenicity in *P. aeruginosa,* then perhaps this enzyme is important across the bacterial kingdom. Furthermore, if L-SD is important for pathogenicity an innate property of prokaryotic L-SD (4FeS), could allow for selective attack on this enzyme while leaving the eukaryotic homolog unhindered (Cicchillo *et al.*, 2004).

In conclusion this study is the first attempt to characterize the highly bioenergetic L-serine deaminase pathway in *P. aeruginosa*. We demonstrate the both *P. aeruginosa* L-SDs, SdaA and SdaB contribute to L-serine deamination. The amino acids glycine and serine were shown to induce L-SD activity *in vivo*. In sharp contrast to *E. coli* L-SD regulation, the amino acid L-leucine was shown to inhibit L-SD activity. Lastly, suicide vectors were constructed which allow for selective mutation of the *sdaA* and *sdaB* genes on any *P. aeruginosa* strain of interest.

Microbiologists have long looked for the “silver bullet” to attack microorganisms. As a result, the bioenergetics of infection has been overlooked as a possible source of therapeutic targets. In this regard, the bioenergetics of infection holds great promise in the dynamic interactions between host and pathogen. Against an ever-evolving microbe, it is imperative to seek alternate modes of attack. *In the absence of energy, there is no life*.

## ACKNOWLEDGEMENT

We acknowledge Dr. Phil Hartman at Texas Christian University for providing the worms and protocols for the *C. elegans* assay. A special thanks to Dr. Dee Mills for arranging the “Nematode Force” meeting. We thank Mathee Crew especially, Dr. Kok-Fai Kong, Dr. Robert Sautter, and the late Robert J. Smiddy for their guidance and insights. We thank Justin Ceballos for formatting the thesis into a manuscript.

